# The multimodal transcriptional response of denervated skeletal muscle involves alterations in cholesterol homeostasis impacting muscle size

**DOI:** 10.1101/2024.09.30.615874

**Authors:** Cristofer Calvo, Casey O. Swoboda, Fabian Montecino Morales, Siddhant Nagar, Michael J. Petrany, Chengyi Sun, Hima Bindu Durumutla, Mattia Quattrocelli, Douglas P. Millay

## Abstract

The development and maintenance of the neuromuscular junction (NMJ) requires reciprocal signals between the nerve terminals and the multinucleated skeletal muscle fiber (myofiber). This interaction leads to highly specialized transcription in the sub-synaptic or NMJ myonuclei within mature myofibers leading to clustering of acetylcholine receptors (AChRs). Here, we utilized single-nucleus RNA sequencing (snRNA-seq) to delineate the transcriptional response of myonuclei to denervation. Through snRNA-seq on skeletal muscle from two independent mouse models of denervation, sciatic nerve transection and amyotrophic lateral sclerosis, we identify a multimodal transcriptional response of NMJ-enriched genes and an alteration in cholesterol homeostasis in both slow and fast myofibers. *Gramd1*, a family of genes involved in non-vesicular cholesterol transport, are enriched at the NMJ in innervated muscle and upregulated in both models of denervation by the NMJ and extrasynaptic myonuclei. *In vivo* gain and loss of function studies indicate that NMJ-enriched *Gramd1 genes* regulate myofiber sizes independent of an obvious impact on AChR clustering. We uncovered a dynamic transcriptional response of myonuclei to denervation and highlight a critical role for cholesterol transport to maintain myofiber sizes.

## Introduction

The neuromuscular junction (NMJ) is a synapse between the nerve terminals of the motor neurons and the plasma membrane of muscle cells (myofibers). During embryonic development, a plethora of factors [1–6] derived from the nerve and the muscle initiate the formation of a highly specialized membrane region in the myofiber that leads to the formation of a cluster of acetylcholine receptors (AChRs). AChR clustering is essential for the transmission of the nerve impulse and, ultimately, contraction of myofibers [7]. Disruption of the NMJ due to motor neuron diseases [8], acute nerve injury [9], or aging [10–12] results in myofiber denervation. Consequently, myofibers begin to atrophy and the AChR clusters become unstable, decrease in size and degenerate if the connection is not re-established with the motor neuron [9, 13].

In the past three decades major factors that are required for the formation, maturation and maintenance of the AChR clusters have been identified. Agrin, a glycoprotein derived from the motor neuron terminals [2], binds to the complex of Lrp4/MuSK on the muscle membrane, causing the dimerization and phosphorylation of MuSK [3]. MuSK activation initiates a downstream signaling cascade mediated by Dok-7 [14, 15] that leads to the clustering of AChRs by the scaffolding protein Rapsyn [6, 16]. Even though AChRs are embedded in the post-synaptic muscle membrane and are localized in cholesterol-rich domains [17], it is unclear how the modulation of the lipid environment of the membrane impacts the AChR clustering and NMJ function.

Multinucleated myofibers compartmentalize the transcription of NMJ genes within a few nuclei in the subsynaptic region [18]. Due to the lack of tools to isolate and study the NMJ nuclei, there is not yet a comprehensive understanding of the transcriptional regulation of NMJ genes. Recently, we and others performed single-nucleus RNA sequencing of skeletal muscle and revealed a series of transcripts with unknown function at the NMJ [19–21]. Loss of specialized transcription at the NMJ is a hallmark of denervation and aging [22–25]. *Musk* and AChR genes are increased in the extrasynaptic compartment of the myofiber after surgical denervation or selective blockade of neuromuscular transmission, but it is unclear whether muscle electrical activity or neural derived factors regulate the synaptic-specific expression of NMJ genes [23, 26]. While major regulators of NMJ development (MuSK, AChR genes) are increased in denervated myofibers, two contrasting responses occur within a given myofiber membrane: 1) degeneration of synaptic AChR clusters and 2) formation of new immature extrasynaptic AChR clusters, suggesting that other factors with compartment-specific transcriptional control may also regulate the AChR clustering responses after denervation [9, 27]. The resolution associated with genomic technologies may help to identify new modalities of the NMJ transcriptional response and novel factors that regulate the response of the NMJ during myofiber denervation, aging or disease.

In this study, we explored the early myonuclear transcriptional response to denervation in two independent mouse models at single nucleus resolution. The response is similar between the two models of denervation and identifies a series of regulated genes with unknown functions in myofibers. We uncover a multimodal transcriptional response of NMJ-enriched genes following denervation including genes that: a) continue to be specifically expressed at the NMJ b) are downregulated in the NMJ myonuclei and are not upregulated extrasynaptically c) lose specialized transcription and are transcribed by extrasynaptic nuclei. The regulation of *Gramd1b* was intriguing because it was one of the few genes that increased in both NMJ and extrasynaptic nuclei after denervation and it is a known regulator of cholesterol distribution. While, there is limited understanding of how cholesterol levels at the NMJ are regulated in homeostasis or denervation, Aster proteins (Aster-a, b and c encoded by *Gramd1a*, *Gramd1b* and *Gramd1c*) are central for the trafficking of accessible cholesterol between membrane compartments (especially plasma membrane) and the endoplasmic reticulum (ER) [28, 29]. Aster proteins are localized in the ER membrane and respond upon increased levels of cholesterol at the plasma membrane (PM) by translocating to the ER-PM contact sites. Thus, Aster proteins have been proposed as ER cholesterol sensors that regulate the homeostasis and distribution of cholesterol in various tissues [30–33]. We show that denervation is associated with increased levels of cholesterol and gain and loss of function experiments *in vivo* revealed that *Gramd1* genes regulate myofiber size. Together, our single-nucleus RNA sequencing in denervated skeletal muscles highlights the role of cholesterol homeostasis in skeletal muscle biology.

## Results

### A conserved myofiber transcriptional response to denervation in adult mice

To elucidate transcriptional responses of myonuclei to neuromuscular junction (NMJ) perturbations, we performed single-nucleus RNA sequencing from the tibialis anterior (TA) muscle of the ALS mouse model SOD1^G93A^ at 2 months of age, which is an early stage of the disease characterized by both denervation and regeneration of nerve-muscle interactions [34]. The transcriptomes of ∼3500 nuclei from the SOD1^G93A^ TA were clustered and compared to a UMAP of snRNAseq from 2 month old wild-type (WT) TA [20] (Supplemental Figure 1A). In the SOD1^G93A^ sample we observed the presence of two myonuclear clusters with gene expression profiles that were not found in the WT TA myonuclear clusters. These two clusters were *Titin*^+^, exhibited low expression of the major myosin genes of the TA muscle, *Myh1* (Type IIx) and *Myh4* (Type IIb), but absence of embryonic *Myh3* (Supplemental Figure 1B), suggesting these myonuclei have been reprogrammed away from the baseline state and are not derived from newly formed myofibers. Through integration of the SOD1^G93A^ mouse dataset with the WT control we observed one myonuclear cluster unique to SOD1^G93A^ muscle (ALS myonuclei) (Figure 1A). The presence of one myonuclear cluster after integration identifies the most divergent transcriptional signature SOD1^G93A^ muscle.

**Figure 1.**
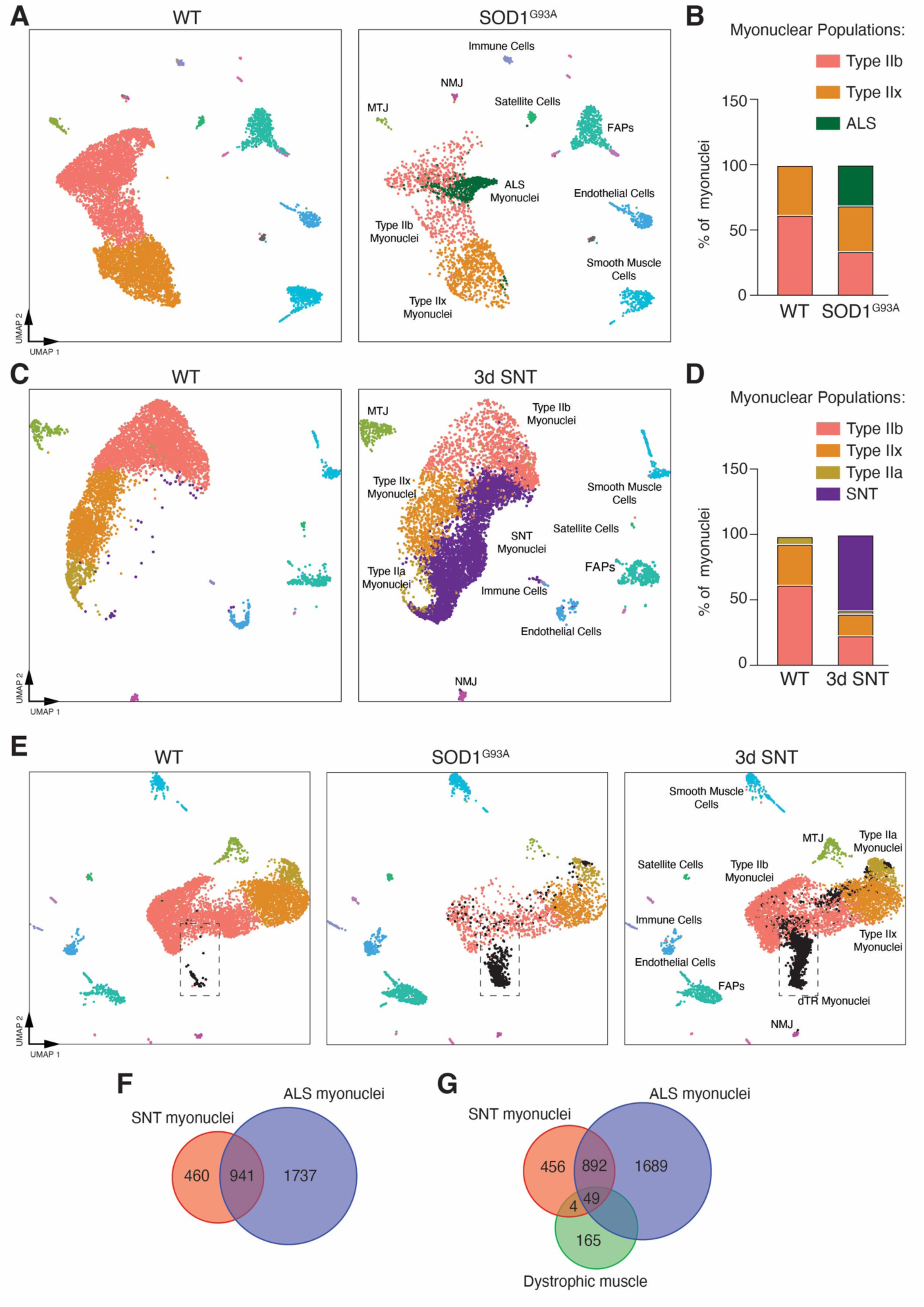
Single nucleus RNA sequencing reveals a conserved myofiber transcriptional response to denervation in adult mice. (**A**) Integrated UMAP split by snRNA-seq data from wild-type (WT) and SOD1^G93A^ tibialis anterior (TA) muscles at 2 months of age. The 10 populations are color-coded and labeled. (**B**) Proportions of myonuclear populations from WT and SOD1^G93A^ snRNA-seq data. (**C**) Integrated UMAP split by snRNA-seq data from TA muscles from adult (5 months of age) WT and 3 days after sciatic nerve transection (3d SNT). The 10 nuclear populations are color-coded and labeled. (**D**) Proportions of myonuclear populations in WT and 3d SNT samples. (**E**) Integrated UMAP split by snRNA-seq data from 5 months WT, SOD1^G93A^ and 3d SNT TA muscles. We refer to the conserved population between SOD1^G93A^ and 3d SNT as denervated transcriptionally reprogrammed (dTR) myonuclei. (**F**) Venn diagram of upregulated genes from SNT myonuclei compared to ALS myonuclei. (**G**) Venn diagram comparing upregulated genes from SNT myonuclei, ALS myonuclei and dystrophic muscle.

When comparing proportions of myonuclei between WT and SOD1^G93A^, the percentage of Type IIx myonuclei remained similar while Type IIb myonuclei were reduced (Figure 1B), suggesting that the ALS-responsive myonuclei emerge within Type IIb myofibers consistent with previous observations that Type IIb myofibers are more susceptible in ALS pathogenesis [35, 36]. Of note, we were unable to identify the Type IIa myonuclei likely due to the relatively low number of nuclei captured in the SOD1^G93A^ mouse. Differential gene expression analysis shows that the ALS-responsive myonuclei has 1156 upregulated and 868 downregulated genes when comparing with IIx and IIb SOD1^G93A^ myonuclei (Supplemental Table 1). Gene ontology analysis of the differentially expressed genes (DEGs) in the ALS myonuclei revealed that the downregulated genes, such as *Tropomyosin* (*Tpm1* and *Tpm2*), *Dystrophin* (*Dmd*) and *Mef2c*, are related with muscle structure, contraction, or development (Supplemental Figure 1C), indicating a partial loss of myonuclear identity. Conversely, the upregulated genes are involved in nerve morphogenesis, synapses and plasma membrane (Supplemental Figure 1C), including known factors for cell-adhesion like *Ncam* and *Igfn1* as well as other transcripts that have not been previously associated with the SOD1^G93A^ mouse model including *Robo2*, *Dlg2*, *Nrcam*, and *Gramd1b*. Through RT-qPCR and smRNA-FISH we validated upregulation of multiple genes identified as enriched in the ALS myonuclei (Supplemental Figure 2, A and B). These data indicate a heterogenous transcriptional response of different myonuclear populations in the SOD1^G93A^ mouse.

To validate that the myofiber transcriptional response we observed in the SOD1^G93A^ mouse model is a general response due to disruption of the connection between nerves and myofibers or specific to the SOD1^G93A^ transgene, we performed snRNA-seq on TA muscles following three days of sciatic nerve transection (3d SNT), a model that results in denervation of all myofibers. We profiled ∼7000 nuclei and after clustering, we observed two clusters (Supplemental Figure 3A) with *Titin* expression, but low *Myh1* and *Myh4* expression (Supplemental Figure 3B). We integrated the 3d SNT dataset with the WT control and found a myonuclear cluster specific to denervated muscle (SNT myonuclei) (Figure 1C) and this population represented 55% of all myonuclear populations in the 3d SNT model (Figure 1D). In the 3d SNT sample, we detected reductions in proportions of Type IIb, Type IIx, and Type IIa myonuclei when compared with the WT control dataset indicating that in the SNT mouse model, unlike the SOD1^G93A^, all fiber types of the TA muscle have a prominent transcriptional response to denervation. DEGs from 3d SNT myonuclei compared with WT myonuclei revealed downregulated genes involved in muscle development, structure and metabolism including *microRNA-133*, *Dystrophin* (*Dmd*), *Creatine Kinase*, *Nr4a1,* and *Sarcoglycan* (*Sgcd*) as well as upregulated genes involved in development, atrophy, and neuromuscular synaptic activity such as *Runx1*, *Musk*, *Scn5a*, *Utrophin* (*Utrn*) and *Myogenin* (*Myog*) (Supplemental Table 2). To determine if the divergent populations observed in the SOD1^G93A^ and SNT models exhibited similar transcriptional identities, we integrated all four snRNAseq datasets: SOD1^G93A^, 3d SNT mouse models, and their respective WT controls. We observed a distinct cluster present in both SNT and SOD1^G93A^ samples, indicating a common transcriptional signature, and we named this population denervated transcriptionally reprogrammed (dTR) myonuclei (Figure 1E).

To better understand the common gene signature within the dTR myonuclei, we compared the most significantly upregulated genes in both models to their respective controls, which revealed an overlap of 941 genes (*Chrng*, *Dlg2, Sema6a, Gramd1b, Musk*) representing a conserved denervation response. We also observed that both SNT and ALS myonuclei possessed a gene signature that was enriched to the specific model (Figure 1F, Supplemental Table 3). For SNT, the enriched gene signature included *Flnc*, *Myh9*, *Sema3f* and *Chrna9* whereas *Robo2*, *Sox11*, *Acta2*, *Tgfb1* and *Gramd1a* were enriched in the ALS model, and the divergent gene signature between models could be explained by complete denervation in SNT in contrast to ongoing denervation and reinnervation in the ALS model.

To determine if these responses were due to a general muscle injury, we also compared the upregulated genes in the SNT and ALS myonuclei to upregulated genes in a mouse model of muscular dystrophy [38], which is a genetic muscle pathology that is not caused by a direct disruption of the nerve-muscle synapse. Here, we detected 49 common genes (Figure 1G), indicating the transcriptional response we observed in 3d SNT and SOD1^G93A^ is not a general response to a muscle insult, but rather denervation-specific. We next asked if the dTR signature is conserved at later timepoints following SNT, which can involve regeneration of the nerve-muscle synapse. We integrated our datasets (3d SNT + SOD1^G93A^) with publicly available snRNAseq performed 14 days following SNT (14d SNT) in the gastrocnemius muscle [39], and also performed qPCR for genes enriched in dTR myonuclei at multiple timepoints after SNT. After integration, the dTR myonuclear cluster is present in the 14d SNT and the 3d SNT + SOD1^G93A^ datasets with similar proportions (Supplemental Figure 4, A and B), suggesting that the dTR myonuclear response is conserved across later time-points after denervation. Of the 941 common genes enriched in dTR myonuclei (Supplemental Table 3), we found known NMJ genes like *Musk* and *Chrng*, but also other factors with unknown function in denervated skeletal muscle such as *Sema6a*, *Dlg2*, *Zfp697*, *Gramd1b,* and *Nrcam*. We validated upregulation of these genes at various timepoints after SNT mouse model including at 28 days when evidence of reinnervation is detected (Supplemental Figure 4C). We found that these genes maintain higher expression levels even after 28 days following SNT (Supplemental Figure 4D), suggesting that these candidate genes could impact remodeling associated with muscle denervation and reinnervation.

### NMJ-enriched genes exhibit a multimodal transcriptional response following denervation

In previous studies [19–21], we and others provided insight into NMJ-enriched genes that were determined by single-nucleus RNA sequencing. It is well known that genes normally enriched at the NMJ, such as *Musk* and AChR family of genes, lose their compartmentalized transcription following denervation [26, 40–42] and are expressed by extrasynaptic nuclei, but it is unclear if all NMJ-enriched genes exhibit this regulation. Using our 3d SNT snRNAseq data, we determined the response of NMJ-enriched genes (total of 128 genes identified from our WT snRNAseq data previously published [20]) to denervation by analyzing expression within the NMJ and extrasynaptic compartments. Heatmap analysis of these 128 NMJ genes highlights a multimodal transcriptional response (Figure 2A) where genes are unchanged between the NMJ and extrasynaptic myonuclei, genes are increased in both compartments, and genes that are reduced at the NMJ. Figure 2B shows comparison of the 128 genes normally enriched at the NMJ to: a) upregulated DEGs found in the NMJ population in the SNT data (35 total, blue) b) downregulated DEGs in the NMJ population in the SNT data (86 total, orange) c) upregulated genes in the dTR myonuclei (820 total, pink), which would represent extrasynaptic expression. Supplemental Table 4 shows genes within each of these categories. 81 NMJ-enriched genes do not respond to denervation (*Pdzrn4*, *Ryr3*, *Ano4*), suggesting the expression of these genes by NMJ myonuclei is not dependent on the presence of the nerve. Indeed, this category includes *Etv5*, an upstream regulator of the NMJ gene program, that does not transcriptionally respond to the lack of innervation during embryonic development or denervation in the adult muscle [43]. 4 genes (*Gramd1b*, *Zfp697*, *Atp13a3, Col13a1*) are increased in both the NMJ and extrasynaptic compartments. We detected the canonical response to denervation where 30 genes (*Musk*, *Lrp4*, *Ncam1*) increased extrasynaptically. We also detected 12 genes (*Ufsp1*, *Ache*, *Chrne*) that are downregulated at the NMJ and 1 gene (*Osbpl10*) that is both downregulated at the NMJ and increased in extrasynaptic compartments. Supplemental Table 4 contains the list of genes in each of these categories. These data indicate a more complex transcriptional response to denervation beyond the concept that NMJ genes simply lose compartment-specific expression, and this multimodal response could be a molecular underpinning of the response of muscle to denervation.

**Figure 2.**
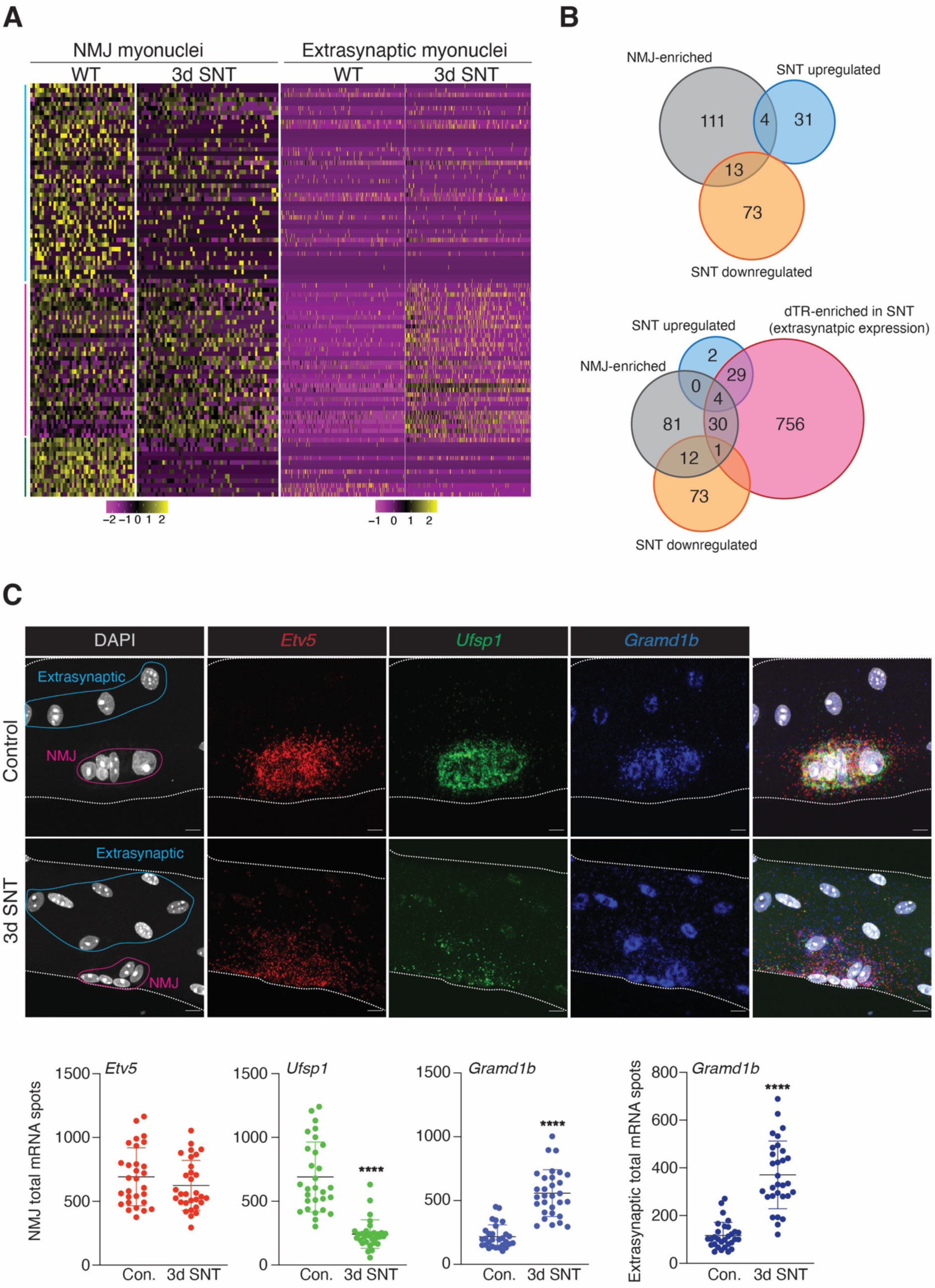
NMJ-enriched genes exhibit a multimodal transcriptional response following denervation. (**A**) Genes normally enriched at the NMJ are represented on a heatmap, comparing their differential expression in NMJ and extrasynaptic myonuclei in innervated WT muscle and after 3d of SNT. Color lines on left of UMAP highlight the three major transcriptional responses: genes are unchanged between the NMJ and extrasynaptic myonuclei (blue), genes are increased in both compartments, and genes that are reduced at the NMJ. (**B**) Venn diagram comparing WT NMJ-enriched genes, 3d SNT NMJ upregulated, and 3d SNT NMJ downregulated (top). Venn diagram comparing WT NMJ-enriched genes, SNT NMJ upregulated, SNT NMJ downregulated, and dTR-enriched genes from the SNT dataset highlighting the multimodal transcriptional response of NMJ genes following SNT (bottom). (**C**) Representative images of smRNA-FISH for *Etv5, Ufsp1*, and *Gramd1b* on EDL myofibers from WT mice undergoing 3d SNT or control conditions. NMJ and extrasynaptic myonuclei are delineated by pink and blue dotted lines, respectively. Quantification of the mRNA spots is shown at the bottom. Each dot represents a single NMJ or extrasnyaptic region (three independent experiments, 9-10 myofibers per experiment). Scale bar, 10μm. Data are presented as mean ± SD. Statistical test used was unpaired t-test; ****p<0.0001.

To validate the NMJ multimodal transcriptional response to denervation we performed a multiplex smRNA-FISH in single myofibers from the extensor digitorum longus [25] following 3d of SNT. We selected *Etv5* (continues to be specifically expressed at the NMJ), *Ufsp1* (downregulated in the NMJ myonuclei), and *Gramd1b* (increased at the NMJ and extrasynaptically). In the control innervated EDL myofibers we observed colocalization and enrichment of all three transcripts at the NMJ (Figure 2C). While *Etv5* and *Ufsp1* were specific to NMJ myonuclei, we observed *Gramd1b* transcripts in extrasynaptic nuclei albeit at lower levels compared to NMJ myonuclei. Following 3d of SNT, we detected decrease of transcripts for *Ufsp1*, no changes for *Etv5* and increase of transcripts for *Gramd1b* in both NMJ and extrasynaptic compartments (Figure 2C). We also validated dynamic changes of *Gramd1b* expression in the SNT model by smRNA-FISH (Supplemental Figure 5A), where we determined that the peak of expression is two weeks after denervation. Since *Gramd1b* is a marker of the dTR myonuclear cluster in both 3d SNT and SOD1^G93A^, and is one of the few detected transcripts that are upregulated in NMJ and extrasynaptic myonuclei, we hypothesized that this factor could be a critical regulator of the physiological response of myofibers to denervation.

### *Gramd1* expression levels are associated with cholesterol alterations following skeletal muscle denervation

*Gramd1b* encodes for the protein Aster-b that functions to regulate cholesterol homeostasis by shuttling this lipid between different cellular compartments [28, 29, 32, 33, 44]. Since *Gramd1b* is highly increased following denervation, we tested if additional genes involved in cholesterol homeostasis are also regulated. *Lldr* is involved in the uptake of cholesterol and does not change during denervation, however we found increased expression of *Srebf-2* and *Hmgcr*, which encode for proteins essential for the biosynthesis of cholesterol. (Figure 3A). *Abca1,* a regulator of cholesterol export, is also increased following SNT (Figure 3A). These data suggest a temporal transcriptional response to remodel cholesterol homeostasis in muscle after denervation. Furthermore, we measured the levels of total cholesterol in the TA (mainly fast myofibers, Type IIx and Type IIb) and also extended this analysis to the soleus (slow myofibers, Type I and Type IIa) muscle to allow a more comprehensive understanding of cholesterol homeostasis across myofiber types. At baseline, we observed elevated levels of total cholesterol in the soleus compared to the TA (Figure 3B). Two weeks following SNT, total cholesterol levels were increased in both TA and soleus muscles compared to the contralateral innervated muscle (Figure 3B). We then determined if cholesterol levels are correlated with the *Gramd1b* mRNA levels and found that *Gramd1b* levels are elevated in the soleus at baseline compared to the TA, and that *Gramd1b* is increased in both muscles after denervation (Figure 3C). *Gramd1a*, a paralog gene that also encodes for Aster proteins, exhibits a similar pattern of gene expression between the soleus and TA, but it is more modestly upregulated following SNT (Figure 3C). These data indicate that the soleus has elevated levels of cholesterol compared to the TA, and that a conserved response to denervation is an increase in cholesterol metabolism.

**Figure 3.**
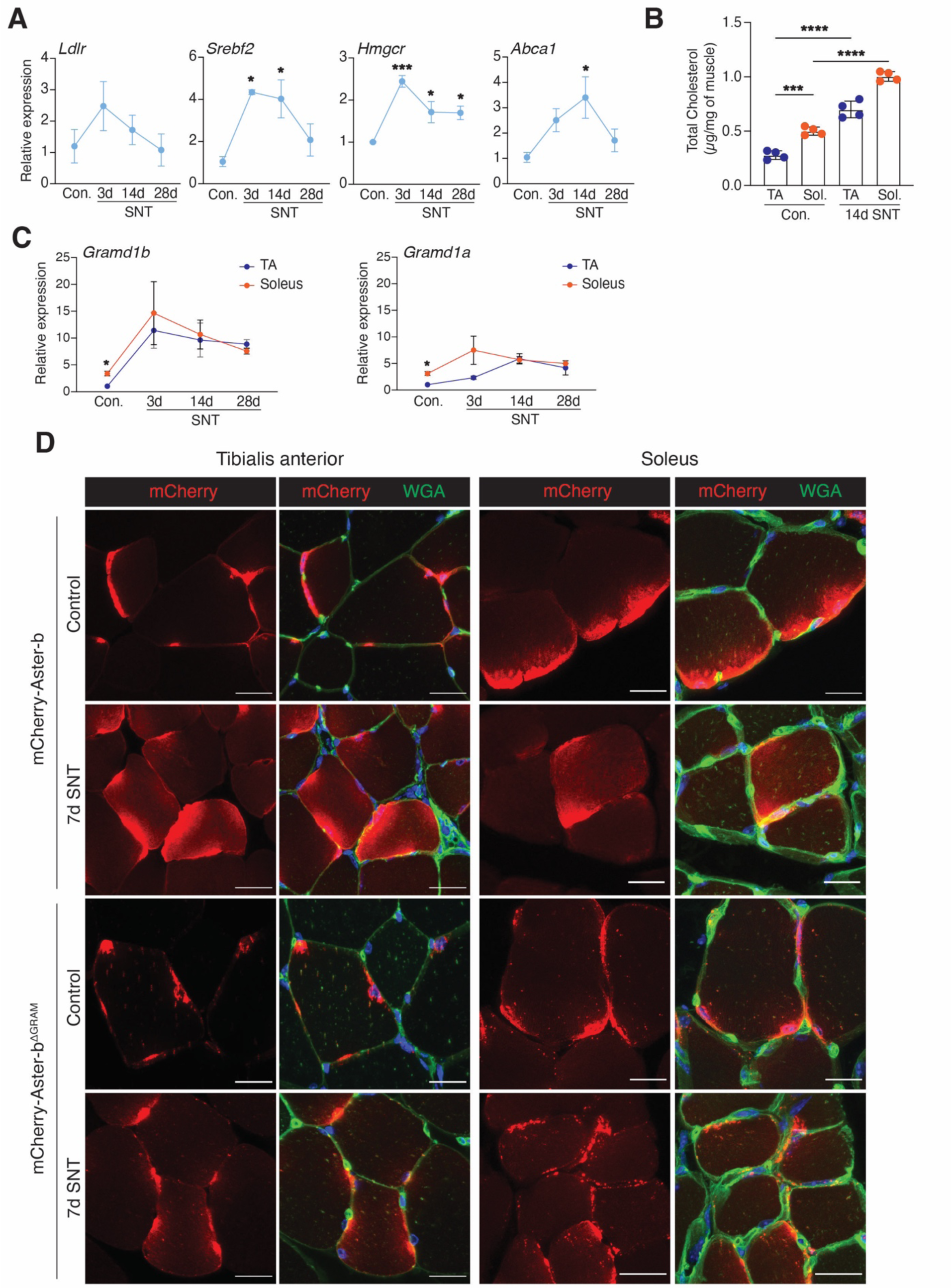
*Gramd1* expression levels are associated with cholesterol alterations in a fiber-type specific manner and following skeletal muscle denervation. (**A**) RT-qPCR analysis for the indicated genes associated with cholesterol metabolism from TA muscles at baseline and different time points following SNT (n=3 per time point). (**B**) Total cholesterol (µg/mg) from the TA and soleus muscles in control and 14d SNT (n=4). (**C**) RT-qPCR analysis of *Gramd1a* and *Gramd1b* comparing the mRNA levels between the TA and soleus in control (TA n=3, Soleus n=4) and SNT at different time points (TA 3d and 28d n=3, TA 17d n=4, Soleus 3d and 17d n=3 and Soleus 28d n=2). (**D**) MyoAAV-mCherry-Aster-b or MyoAAV-mCherry-Aster-b^ΔGRAM^ were injected intramuscularly in the hindlimbs of WT mice and mice were subjected to unilateral SNT for 7 days. Representative images of cross sections of TA and soleus muscles comparing localization of Aster-b and Aster-b^ΔGRAM^ between control and 7d SNT. This experiment was repeated three times. Scale bars, 10μm. Data are presented as mean ± SD. Statistical test used were (**A**) and (**B**) one-way ANOVA with Tukey’s correction for multiple comparisons; (**C**) multiple unpaired t-tests; *p<0.05, **p<0.01, ***p<0.001, ****p<0.0001.

Since Aster-b regulates the transport of the accessible cholesterol pool from the plasma membrane (PM) to endoplasmic reticulum (ER) [28, 29, 32], we tested if the increase of total cholesterol we observed after denervation translates to a change of the accessible cholesterol pool at the skeletal muscle membrane. We generated two Aster-b constructs with mCherry epitope tags as sensors for the accessible cholesterol pool in skeletal muscle, where one contained full-length Aster-b including the GRAM domain that is needed for cholesterol binding and the other construct contains Aster-b that lacks the GRAM domain (Aster-b^ΔGRAM^) [28, 29]. We delivered MyoAAV-mCherry-Aster-b into hindlimbs of WT mice, performed unilateral SNT, and harvested muscles 7 days after SNT. In the TA and soleus muscles at baseline, we observed perinuclear and sarcolemma localization of mCherry-Aster-b, whereas in the denervated muscles we observed diffusion of the protein inside of the myofiber indicating there is redistribution of cholesterol after denervation (Figure 3D). As a control, we performed a similar experiment with MyoAAV-mCherry-Aster-b^ΔGRAM^ and did not detect changes in localization of mCherry between the control and denervated muscles (Figure 3D), indicating that the mCherry-Aster-b is faithfully detecting accessible cholesterol. We conclude that the accessible cholesterol pool is increased in the myofiber membrane system following denervation in both TA and soleus muscles, and Aster-b responds to the cholesterol gradient to mobilize this lipid between myofiber compartments.

### Genetic manipulations of *Gramd1* genes alters myofiber sizes

While genetic deletion of *Gramd1b* in the mouse has been reported to not compromise viability *in vivo* [28, 31], in our colony we observed reduced numbers of *Gramd1b*^-/-^ mice upon genotyping (postnatal day 7-10) (Supplemental Figure 5B). Since we observed normal NMJ development through assessment of AChRs and nerves in the diaphragm of E18 WT and *Gramd1b*^-/-^ animals (Supplemental Figure 5C) regardless of their likelihood to survive, we conclude that the lethality is not due a disruption of the NMJ. Therefore, *Gramd1b* is dispensable for NMJ development.

We also considered the role of *Gramd1b* in denervated myofibers since cholesterol levels are increased after denervation (Figure 3C) and the expression of *Gramd1b* increases in both the NMJ and extrasynaptic compartments (Figure 2C). To recapitulate the denervation transcriptional response of elevated *Gramd1b* in a muscle-specific manner, we delivered AAV9-Gramd1b, or AAV9-GFP as a control, to TA muscles of adult WT mice and analyzed innervated muscles after 28 days. In TAs injected with AAV9-Gramd1b, we observed overexpression of *Gramd1b* in myofibers (Figure 4A), reductions in myofiber CSA (Figure 4A), and reduced AChR cluster areas (Figure B) with no impact on overall morphology of AChRs or innervation. These data indicate that upregulation of *Gramd1b* decreases myofiber sizes and impacts regulation of AChR cluster sizes.

**Figure 4.**
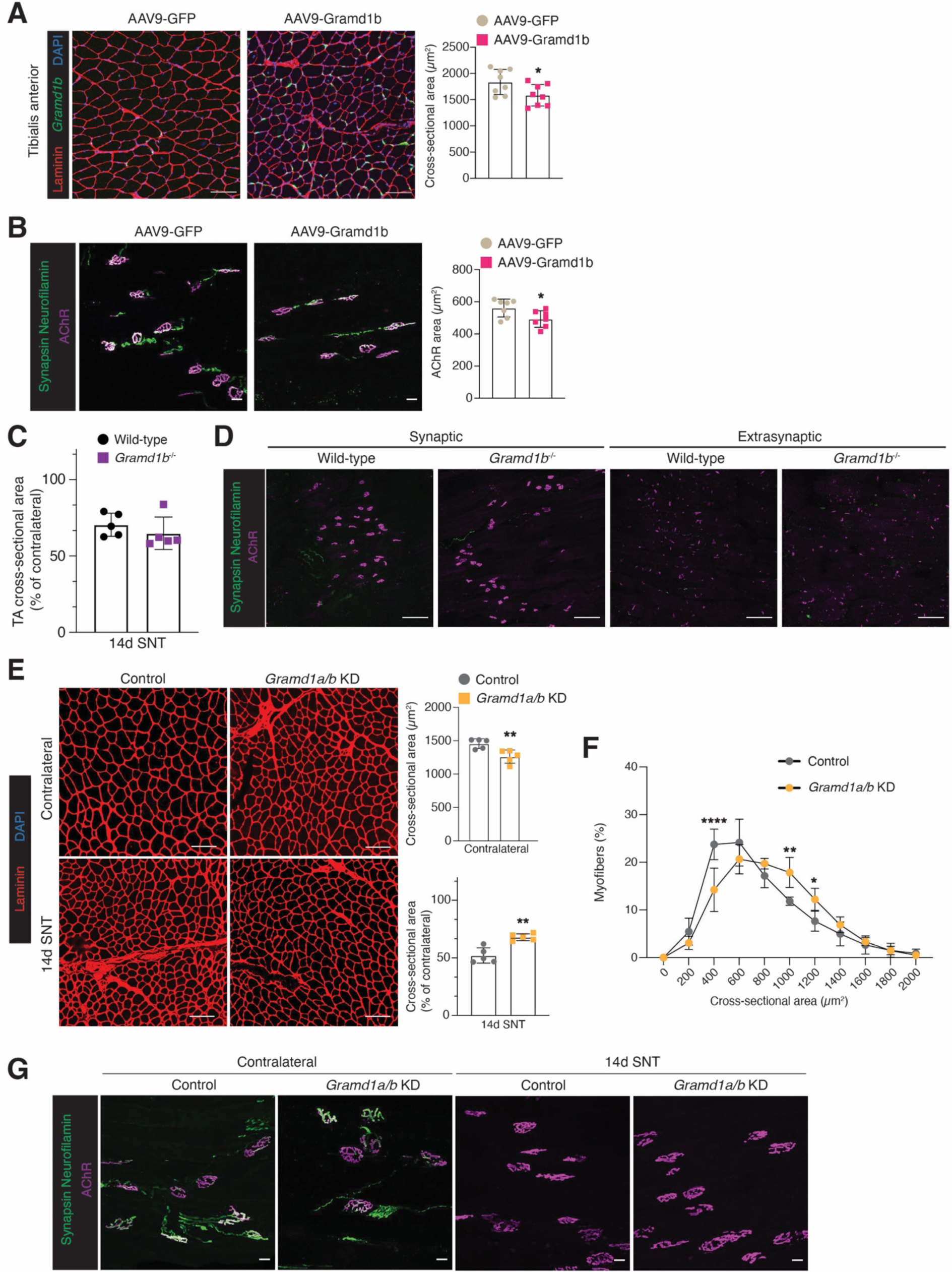
Regulation of myofiber sizes by genetic manipulation of *Gramd1* genes. (**A**) Representative staining of smRNA-FISH for *Gramd1b* in cross sections of TA muscles that were injected with AAV9-GFP or AAV9-Gramd1b for 28 days. Quantification of cross-sectional area (CSA) is shown on the right. Scale bars, 100μm (n=8). (**B**) Representative staining of AChRs, axons (neurofilament), and nerve terminals (synapsin) of TA muscles after one month of AAV mediated overexpression of *Gramd1b*. Quantification of AChR area (µm^2^) is shown on the right side. Scale bars, 10μm. (**C**) Percentage (%) of contralateral CSA of WT and *Gramd1b*^-/-^ TA muscles 14 days following SNT. (**D**) Representative staining of AChRs, axons (neurofilament) and nerve terminals (synapsin) of synaptic and extrasynaptic areas from WT and *Gramd1b*^-/-^ gastrocnemius muscles. Scale bars, 100μm. (**E**) Representative laminin-stained TA cross sections of *Gramd1a/b* KD mice one month after delivery of gRNAs via intramuscular injections into *Rosa26*^Cas9^ mice. Cross sections of contralateral and denervated muscles are shown. Quantification of cross-sectional area (CSA) of the contralateral muscle or percentage (%) of contralateral CSA of control and *Gramd1a/b KD* TA muscles two weeks following SNT (n=5) is shown on the right. Scale bars, 100μm. (**F**) CSA distribution of control and *Gramd1a/b KD* TA muscles two weeks following SNT (n=5). (**G**) Representative staining of AChRs, axons (neurofilament), and nerve terminals (synapsin) of control and *Gramd1a/b KD* TA muscles. Scale bars, 10μm. Data are presented as mean ± SD. Statistical tests used were (**A**), (**B**), (**C**), and (**E**) unpaired t-test; (**F**) two-way ANOVA with Tukey’s correction for multiple comparisons; *p<0.05, **p<0.05, ****p<0.0001.

Next, we asked whether loss of *Gramd1b* can also influence the myofiber denervation response. The atrophy response of the TA of *Gramd1b^-/-^* mice was as pronounced as control animals, based on percentage of contralateral CSA (Figure 4C). Deletion of *Gramd1b* also did not impact the sizes of synaptic or extrasynaptic AChR clusters in gastrocnemius muscles four weeks following SNT (Figure 4D). These results suggest that loss of *Gramd1b* does not alter the myofiber denervation response. However, the presence of additional *Gramd1* genes (*Gramd1a*, *b* and *c*) could undermine the effect of loss of only *Gramd1b* in cholesterol regulation. When *Gramd1b* is the main paralog gene expressed, such as in the adrenal glands and ovaries, genetic deletion is sufficient to alter cholesterol homeostasis [28, 31]. In contrast, in the liver and the intestine, two *Gramd1* genes are highly expressed and inactivation of both genes is required to observe a phenotype [30, 33]. In skeletal muscle, we did not detect *Gramd1c* expression (not shown) but showed expression of *Gramd1a* and *Gramd1b* in soleus and TA, and demonstrated that both *Gramd1a* and Gramd1*b* are responsive to denervation (Figure 3C). While *Gramd1a* in gastrocnemius muscles of *Gramd1b*^-/-^ mice is not altered in control or denervated muscles (Supplemental Figure 5D), the presence of this gene could reduce the sensitivity of loss of the other family member on cholesterol homeostasis.

We used a CRISPR/Cas9 mouse model to conditionally inactivate both *Gramd1a* and *Gramd1b* in adult TA muscles [45, 46]. MyoAAV [47] containing gRNAs to target exons 2-8 of both *Gramd1a* and *Gramd1b* (Supplemental Figure 6A) were delivered through intramuscular injections (IM) into the TA/EDL muscles of *Rosa26*^Cas9^ mice [46]. The levels of *Gramd1a/b* were decreased in EDL muscles from the mice that received *Gramd1a/b* gRNAs (*Gramd1a/b* KD) (Supplemental Figure 6B). In the TA, we observed reduced myofiber CSA in the contralateral muscle from *Gramd1a/b* KD mice (Figure 4E). While loss of *Gramd1a/b* in the TA resulted in reduced myofiber CSA at baseline, after denervation we observed an increase of myofiber sizes in *Gramd1a/b* KD mice compared to controls when CSA was normalized to the contralateral (Figure 4E). These data indicate that atrophy is not exacerbated in *Gramd1a/b* KD mice after denervation, which might be expected given the reduced size at baseline. Moreover, a higher percentage of larger myofiber sizes were observed in *Gramd1a/b* KD mice after denervation compared to controls suggesting a mitigation of atrophy after loss of *Gramd1a/b* (Figure 4F). Assessment of AChRs in control and *Gramd1a/b* KD mice, revealed no detectable differences in AChR cluster sizes between genotypes in innervated and denervated muscles (Figure 4G). Overall, these data establish that *Gramd1* genes can impact myofiber sizes, but our data suggest that this size regulation is not caused by a major morphological defect in AChR clustering. It is interesting to note that an increase or decrease of *Gramd1* genes in myofibers results in reduced myofiber sizes. We speculate that precise regulation of *Gramd1* genes and subsequent cholesterol localization must be maintained at a critical threshold and alterations above or below that range can impact muscle size.

## Discussion

In this study, we explored the early transcriptional response of myonuclei to skeletal muscle denervation at a single-nucleus resolution. We were particularly interested in transcriptional changes at the NMJ and the extrasynaptic compartment of myofibers before the muscle exhibits major phenotypic changes such as atrophy, fiber type switching [27, 48] and fibrosis [49] that are not directly due to lack of nerve-muscle communication. These data provide a comprehensive resource of gene expression changes that may be useful to the scientific community to study new pathways involved in skeletal muscle denervation. We identified a dTR myonuclear cluster, which has a largely conserved transcriptional signature in three independent models of denervation. Another recent study also highlighted the heterogeneous myonuclear response to denervation-dependent muscle atrophy and found marker genes (*Runx1, Dlg2*, *Robo2*) that we also observed enriched in the dTR cluster [50], supporting the robustness of this myonuclear transcriptional response.

Even though our previous work did not detect obvious transcriptional heterogeneity beyond myofiber types, NMJ, and MTJ, in adult WT muscle during homeostasis, we and others [20, 22, 25, 51] observed divergent myonuclear populations with a denervation transcriptional signature in mouse and human aged muscle. This divergent transcriptional signature leads to the question of how this response is spatially coordinated in the muscle. One possibility is that certain myofiber types transcriptionally respond faster to the lack of innervation. For example, in our 3d-SNT model we observe almost complete absence of *Myh1* (type IIx myofibers) but not of *Myh4* (type IIb myofibers), suggesting type IIx myofibers lose their identity more quickly. Another possibility might be that the syncytium recruits only a portion of the myonuclei to response to lack of innervation. Pairing our datasets with other genomic approaches such as spatial transcriptomics and ATACseq may provide a better understanding of how the myofiber transcriptionally coordinates the response to denervation.

The resolution of our dataset revealed a multimodal transcriptional response of NMJ genes and expands the current view that NMJ genes only lose transcriptional compartmentalization after denervation [23, 40–42]. *Musk,* a master regulator of NMJ formation, loses transcriptional compartmentalization after denervation, resembling *Musk* expression during embryonic development. Therefore, we speculate that the multimodal transcriptional response of NMJ genes may be a reversal to an embryonic-like state of the myofiber to establish the optimal conditions for reinnervation. Our current understanding is that most NMJ components on myofibers remained localized for weeks after denervation, thus, axons are capable to regenerate at original synaptic sites [52, 53], but also immature AChR clusters form in the extrasynaptic region of the myofiber. The multimodal regulation may be correlated with gene function following denervation where transcripts that retain sub-synaptic expression may stabilize the NMJ structure whereas transcripts that increase in the extrasynaptic compartment may promote degeneration of the NMJ at the synaptic region and formation of extrasynaptic AChR clusters. Moreover, our data revealed 12 NMJ transcripts that exhibit downregulation in denervated muscle, including genes such as *Colq*, *Ache* and *Chrne*, known to be involved in neuromuscular transmission of mature skeletal muscle [7, 54], indicating that electrical activity or neurotrophic factors are required for maintaining the expression of the downregulated transcripts. We previously found that *Ufsp1*, another transcript of this mode of regulation, negatively impacts AChR clustering [20] a phenomenon that has not been well studied at the NMJ. It is possible that the genes downregulated at the NMJ exhibit novel functions that have not been previously explored.

Our snRNA-seq of denervated muscle revealed alterations in genes associated with cholesterol regulation, which has been extensively studied in tissues that rely heavily on cholesterol metabolism and transport such as adipocytes and liver. Here, we found slow myofibers exhibit higher cholesterol levels than fast myofibers at baseline, and that cholesterol is increased following denervation in both muscle types. This is consistent with other studies in rats where cholesterol levels were higher in slow compared with fast muscles [55, 56], and an increase of total cholesterol was detected in the denervated plantaris and gastrocnemius [57]. However, the consequence of altered cholesterol homeostasis on muscle at baseline and after denervation had not been investigated prior to our work here. We found that *Gramd1b*, a factor that transports accessible cholesterol between membrane compartments in different cell types [28, 30, 31, 33], was a component of the multimodal response to denervation where *Gramd1b* is enriched at the NMJ at baseline and its expression is increased in both NMJ and extrasynaptyic compartments after denervation. Through genetic manipulation of *Gramd1* genes, we show that increase of these genes in denervated myofibers may have a deleterious effect on size. However, none of the genetic manipulations caused a major morphological change of denervated NMJs, suggesting that changes in myofiber sizes are likely independent of the NMJ.

The conditional inactivation of *Gramd1a/b* in adult innervated muscles, where both genes are enriched at the NMJ, exhibit decrease of myofiber sizes, contrasting the response we observe in denervated *Gramd1a/b KD* muscles. This phenotype cannot be explained by alterations of the NMJ because no changes in AChR clusters or innervation were observed in *Gramd1a/b* KD mice. There are many possibilities for these divergent results including that *Gramd1* could influence myofibers at NMJ and extrasynaptic compartments and it is difficult to parse apart these effects with our current tools. Myofiber cholesterol levels are also increased in denervated muscles and these proteins respond to cholesterol gradients, therefore, the effect of altered cholesterol transport in myofiber membranes may be different in innervated and denervated muscles. Also, cholesterol has pleiotropic functions in all cell types [58–60] and *Gramd1* genes can impact accessible cholesterol levels within all cellular membrane compartments. Thus, the pathways leading to changes of myofiber sizes in different genetic manipulations and innervation contexts may not be related. For instance, cholesterol transport could impact the plasma membrane and function of signaling receptors or ion channels, but also alter the sarcoplasmic reticulum and the function of calcium handling proteins, and each of these effects could lead to changes in myofiber sizes.

Overall, we identify a complex transcriptional response of myonuclei to denervation involving an upregulation of cholesterol metabolism. We report that cholesterol transport is a critical component of muscle size regulation, although due to technical challenges it is not possible to define how transport is altered. This work lays the foundation for new research avenues to study the consequences of cholesterol metabolism on muscle during various pathologies including denervation and statin-induced muscle weakness.

## Methods

### Sex as a biological variable

Our study examined both male and female mice. We did not analyze the sexes independently.

### Animal care and procedures

Wild-type (WT) control mice used in this study were in the C57BL/6 genetic background. *Gramd1b*^-/-^ mice were a generous gift from Dr. Peter Tontonoz (UCLA) [28]. Briefly, this line was generated on a C57BL/6N background using the CRISPR/Cas9 system targeting exon 7 that leads to an early stop codon. The SOD1^G93A^ mouse was generated as previously described [61] and maintained by Jackson Laboratories (strain #004435). *Rosa26*^Cas9^ mice were generated as described previously [46] and they were crossed with the β-actin-Cre mice to excise the floxed STOP cassette. These mice were maintained by Jackson Laboratories in the C57BL/6J background (strain #026179). Mice were housed at 22.2 °C with humidity of 30–70% and a 14 h light/10 h dark cycle and they were fed *ad libitum*. Experimental mice were sacrificed at ages 3-8 months unless otherwise specified for histological, lipid, and gene expression analyses.

Unilateral sciatic nerve transection (SNT) was performed by transecting a piece of the right sciatic nerve in the mid-thigh region after the mouse was anesthetized with isoflurane. For long-term denervation experiments 5-8 mm were transected, but for acute denervation experiments only 1-2 mm. For nerve crush, sciatic nerve was exposed and pinched for 10 seconds using fine forceps. Muscles were harvested at different time points following SNT or nerve crush and the contralateral muscles were used as control.

### Muscle collection and preparation

To collect muscles for analysis, mice were anesthetized using isoflurane and then cervical dislocation was performed. Muscles were harvested, weighed, and then either flash frozen in liquid nitrogen for molecular analysis, or embedded in tragacanth and frozen in liquid nitrogen-chilled 2-methylbutane for downstream histological analysis. Some cohorts of muscle (both control and experimental) were collected and fixed in 4% paraformaldehyde (PFA)/PBS for 10 minutes, then cryoprotected in 20% sucrose/PBS overnight. Muscles were then embedded in OCT and frozen in liquid nitrogen-chilled 2-methylbutane prior to sectioning. All muscles used for cross section histology were cryosectioned at 10 μm thickness at the muscle mid-belly and for longitudinal histology at 50 μm thickness at various levels of the muscle.

### Cloning, AAV production and delivery

*Gramd1b* cDNA sequence was obtained from the NCBI (NM_001357620.1) and synthesized as gblock from IDT. The genetic construct was cloned into the EcoRI site of the pAAV-MCS (Cell Biolabs, catalog VPK-410) plasmid using In-Fusion cloning (Takara Bio). The MyoAAV-mCherry-Aster-b and the MyoAAV-mCherry-Aster-b^ΔGRAM^ genetic constructs were generated by overlap extension PCR using the gblock as template. The 7x sgRNAs targeting *Gramd1a*/b were selected using the tool CRISPOR v5.2 [62] and assembled as previously described using the multiplex CRISPR/Cas9 assembly kit [63] (Addgene kit # 1000000055). sgRNAs sequences and oligos used for cloning are provided in Supplemental Table 5. This genetic construct containing 7x hU6-sgRNAs was cloned into the pAAV-MCS plasmid after removing the CMV promoter using EcoRI and MluI.

All the AAVs were produced in-house using the MyoAAV 1A or AAV9 capsid as previously described [47, 64]. Briefly, AAV Pro 293T cells (Takara) were plated in 15 cm dishes at a density of 2 x10^7^ cells/dish. The next day, each plate was transfected with 10.5 mg helper plasmid, 5.25 mg of the MyoAAV-1A or AAV9 Rep/Cap plasmid, and 5.25 mg of the gene transfer plasmid. Recombinant virus was harvested from the cells and media, and purified by ultracentrifuge using an iodixanol gradient. AAV titers were quantified by qPCR with C1000 Touch Thermal Cycler (Bio-Rad Laboratories). AAVs expressing GFP or mcherry were used as controls. For delivery, viruses were delivered via intramuscular (IM) injections into the hindlimb muscles using 5x10^10^-1x10^11^ viral genomes (vg) for overexpression experiments or 2.5x10^11^ vg for sgRNAs delivery.

### Immunofluorescence and cross-sectional area (CSA) analysis

Muscle sections were first fixed in 1% PFA for 2 minutes, washed in PBS, permeabilized in 0.2% Triton X-100/1x PBS for 10 minutes, washed in PBS, and blocked in 1% BSA/1% heat inactivated goat serum/0.025% Tween-20/1x PBS for 1 h at room temperature in a humidity chamber. Next, slides were incubated in rabbit anti-laminin primary antibody (1:500; Sigma L9393) for 1 h at room temperature, or overnight at 4 °C, in a humidity chamber. After incubation in primary antibody, slides were washed in PBS, and incubated in goat anti-rabbit Alexa Fluor secondary antibodies (1:1000; Invitrogen) for 30–60 min at room temperature in a humidity chamber. Slides were then washed with PBS and mounted with VectaShield containing DAPI (Vector Laboratories). After immunostaining, representative images were taken on a Nikon A1R confocal microscope for use in CSA analysis. The CSA of individual myofibers were measured using NIS Elements software (Nikon), which can quantify the area within laminin-labeled myofibers.

### Immunofluorescence and Neuromuscular Junction (NMJ) analysis

Muscle sections were permeabilized with 1x PBS 0.5% Triton x-100 for 20-30 minutes and block with 2.5% BSA, 2.5% Goat Serum in 0.5% Triton 1x PBS for 1 h. Primary antibodies (Synapsin-1 (D12G5) XP^®^ Rabbit mAb #5297 and Neurofilament-L (C28E10) Rabbit mAb #2837, Cell Signalling) were incubated overnight at dilution 1:500 in blocking buffer. Next, slides were washed 2x with 1x PBS and incubated in goat anti-rabbit Alexa Fluor secondary antibodies (1:1000; Invitrogen) and Invitrogen™ Alpha-Bungarotoxin (1:1000) for 1 h. After immunostaining, six images were taken per muscle in the 40x objective on a Nikon A1R confocal microscope. Analysis of AChR size was done in the Imaris 10.1 (Oxford Instruments). At least 15 individual AChR clusters were analyzed per animal. Quantification of innervation was done manually using Image J.

### smRNA-FISH on cryosections and EDL myofibers

Single-molecule FISH experiments were performed using RNAscope (ACDBio) following the manufacturer’s protocols. We used 10µm fresh-frozen cross-sections of muscles. Following the completion of the RNAscope fluorescent assay, an immunostaining step was carried out to label myofibers. Sections were blocked for 30 minutes using 1% BSA, 1% heat-inactivated goat serum, and 0.025% Tween-20/1x PBS, followed by incubation for 30 minutes at room temperature with anti-laminin (1:500). Secondary AlexaFluor antibodies (1:1000) (Invitrogen) were then applied at room temperature for 1 h. Sections were mounted using VectaShield with DAPI (Vector Laboratories).

For isolation of Extensor digitorum longus [25] myofibers we used the protocol described previously by Kann and Krauss [65]. Briefly, EDL was harvested and incubated in high-glucose DMEM (HyClone Laboratories) containing 0.2% collagenase type I (Sigma-Aldrich) at 37°C in a cell culture incubator. After a maximum of 1 h of incubation, muscles were triturated until myofibers detached from the tissue. Isolated myofibers were collected, fixed in 4% PFA/1x PBS for 20–30 minutes at room temperature, rinsed in 1x PBS and transferred to 100% methanol, stored at - 20°C until needed. For rehydration, slides were washed in a descending methanol/1x PBS+0.1% Tween-20 (PBST) series (50% MeOH/50% 1x PBST; 30% MeOH/70% 1x PBST; 100% 1x PBST) for 5 minutes each. All slides underwent a 10 minutes hydrogen peroxide incubation, followed by a 10 minutes Protease III incubation at room temperature. Slides were washed with 1x PBST for 5 minutes between incubations. Selected probes were used to hybridize slides at 40°C overnight. The ACDBio user manual provided guidance for the remaining AMP hybridization and HRP incubation procedures.

### Total cholesterol assay

Whole muscles were weighed and washed in ice cold 1x PBS to remove any traces of blood. Muscles were cut into small pieces in a 2 mL microcentrifuge tube. Samples were homogenized in 100 µL of Assay Buffer II/Cholesterol Assay Buffer (Abcam, Cat # ab65390, Cambridge, MA, USA) using a Dounce homogenizer with 12 passes. Homogenate was centrifuged for 10 minutes at 4°C at 13,000 x*g* using cold microcentrifuge to remove any insoluble material and supernatant, containing the total cholesterol fraction, was collected Samples were diluted 1:3 in Assay Buffer II. For standard preparation and plate loading, steps were followed according to the manufacturer’s instructions. Absorbance was measured on a microplate reader at OD 570 nm.

### Single nucleus RNA sequencing (snRNAseq)

Tibialis anterior (TA) muscles from SOD1^G93A^ or C57BL/6 mice were harvested immediately following euthanasia, minced, and placed in homogenization buffer (0.25 M sucrose and 1% BSA in Mg^2+^-free, Ca^2+^-free, RNase-free 1x PBS). Nuclei isolation was done as previously described [20]. After filtration via a 40 μm strainer, nuclei were labeled with Hoechst dye and 0.2 U/μL Protector RNase inhibitor (Roche). All Hoechst stained myonuclei were purified using FACS (BD Aria, 70 μm nozzle) and gathered in sorting buffer containing Protector RNase inhibitor (0.2 U/μL). The nuclei were counted with a hemocytometer, and the concentration was fine-tuned as needed to reach the optimal range for the 10X Chromium chip. The 10X Chromium system was then used to load the nuclei using the Single Cell 3′ Reagent Kit v3.1, following the guidelines provided by the manufacturer. Around 12,000 nuclei were loaded for each operation. Sequencing was performed on an Illumina NovaSeq 6000 System.

### snRNAseq data processing and analysis

Initial read alignment and quantification of FASTQ files were generated using CellRanger/v5.0.1. For each dataset, we corrected for ambient background RNA using CellBender [66]. Reads from CellBender were then imported into Seurat objects (Seurat/v5.1.0) [67], and nuclei with less than 200 unique features or greater than 15% reads mapping to the mitochondrial genome were removed from downstream analysis. Additionally, only features expressed in at least 5 cells from each dataset were retained for downstream analysis. Seurat objects with the remaining nuclei then underwent doublet identification using Solo/v0.2 [68]. Nuclei having higher than the 95th percentile of number of unique features, number of UMIs, and mitochondrial reads were excluded. An additional filtration of any nuclei containing higher than 5% reads mapping to the mitochondrial genome were excluded from all datasets after doublet detection. Also, outlier nuclei were determined by having the top 1% detected UMI counts or unique features sequenced for each dataset and were removed prior to downstream processing. Datasets were normalized using the NormalizeData() function. 2000 variable features for downstream Scaling and PCA analysis were defined using the FindVariableFeatures() function. Data were scaled using the ScaleData() function, and principal component analysis on the variable features was performed using the RunPCA() function. For single dataset processing, initial dimensionality reduction was performed using 20 principal components for the RunUMAP() and FindNeighbors() function. Clustering was performed using the FindClusters() function, with a resolution of 1. Nuclei predicted to be doublets were then subset and then an initial round of dimensionality reduction with the same parameters was performed. If a cluster identified in the first round of dimensionality reduction contained more than 50% of nuclei predicted to be doublets, the cluster was subset prior to rerunning dimensionality reduction functions.

### Gene expression analysis

Total RNA was extracted from muscles samples using Trizol and the RNeasy miniKit (Qiagen) following the manufacturer instructions. cDNA was synthesized using the High-Capacity cDNA Reverse Transcription Kit (ThermoFisher) using random hexamers and 1-2μg of total RNA.

Primers were obtained from OriGene Technologies Inc and commercially synthesized (Custom DNA Oligos; IDT). qPCR was carried out on Biorad CFX 96 using the PowerUp SYBR Green Master Mix (Applied Byosistems), reaction volumes of 20 μL, primer concentration of 400 nM and cDNA concentration of 10ng. Data normalization was done by the 2^−ΔΔCt^ method using the geometric average of two reference genes as described previously [69] . Primer sequences are listed in Supplemental Table 5.

### Statistical Analysis

We used the FDR <0.05 criterion for differentially expressed genes (Upregulated and downregulated genes). Gene ontology analysis was performed using ToppGene (https://toppgene.cchmc.org) using the FDR ≤ E-50 criterion [70]. Sample sizes are noted in the figure legends. Data were processed using GraphPad Prism 9 software. In all graphs, error bars indicate standard deviation (SD). Data were compared between groups using various statistical tests, indicated in the figure legends, and based on number of groups, normality of the data, and variance of standard deviations. The criterion for statistical significance was *p<0.05, **p<0.01,***p<0.001, ****p<0.0001.

### Study approval

Animal studies were conducted in accordance with guidelines from the National Institutes of Health and the Division of Veterinary Services (DVS) at Cincinnati Children’s Hospital Medical Center. The IACUC protocol number for this study was # 2023–0017.

### Data and code availability

Raw sequencing data is in process of being deposited in GEO. Data analysis is described in the Methods. Scripts are available upon request.

## Acknowledgements

We thank members of the Millay laboratory and Vikram Prasad (Molkentin lab) for discussion. We thank Maureen Bender, Scott Blair, Fiona Rowan, and Leah Focke for technical assistance. We would like to acknowledge the assistance of the Research Flow Cytometry Core, Single Cell Genomics Core, DNA sequencing Core and Bio-Imaging and Analysis Facility at Cincinnati Children’s Hospital Medical Center. We thank the laboratory of Dr. Peter Tontonoz (University of California, Los Angeles) for the *Gramd1b*^-/-^ mice. Work in the Millay laboratory is funded by grants to D.P.M. from Children’s Hospital Research Foundation, National Institutes of Health (R01AR068286, R01AG082697). Work in the Quattrocelli laboratory is funded by grants to M.Q. from Children’s Hospital Research Foundation, National Institutes of Health (R01HL166356-01, R03DK130908-01A1, R01AG078174-01).

## Author contributions

C.C. and D.P.M. conceived the project and designed experiments. C.O.S performed the bioinformatic analyses. C.C., F.M.M., S.N., M.J.P., C.S, H.B.D. conducted experiments and analyzed the data. D.P.M. supervised the project. C.C. and D.P.M. wrote the manuscript with input from all authors.

---

**Supplemental Figure 1.**
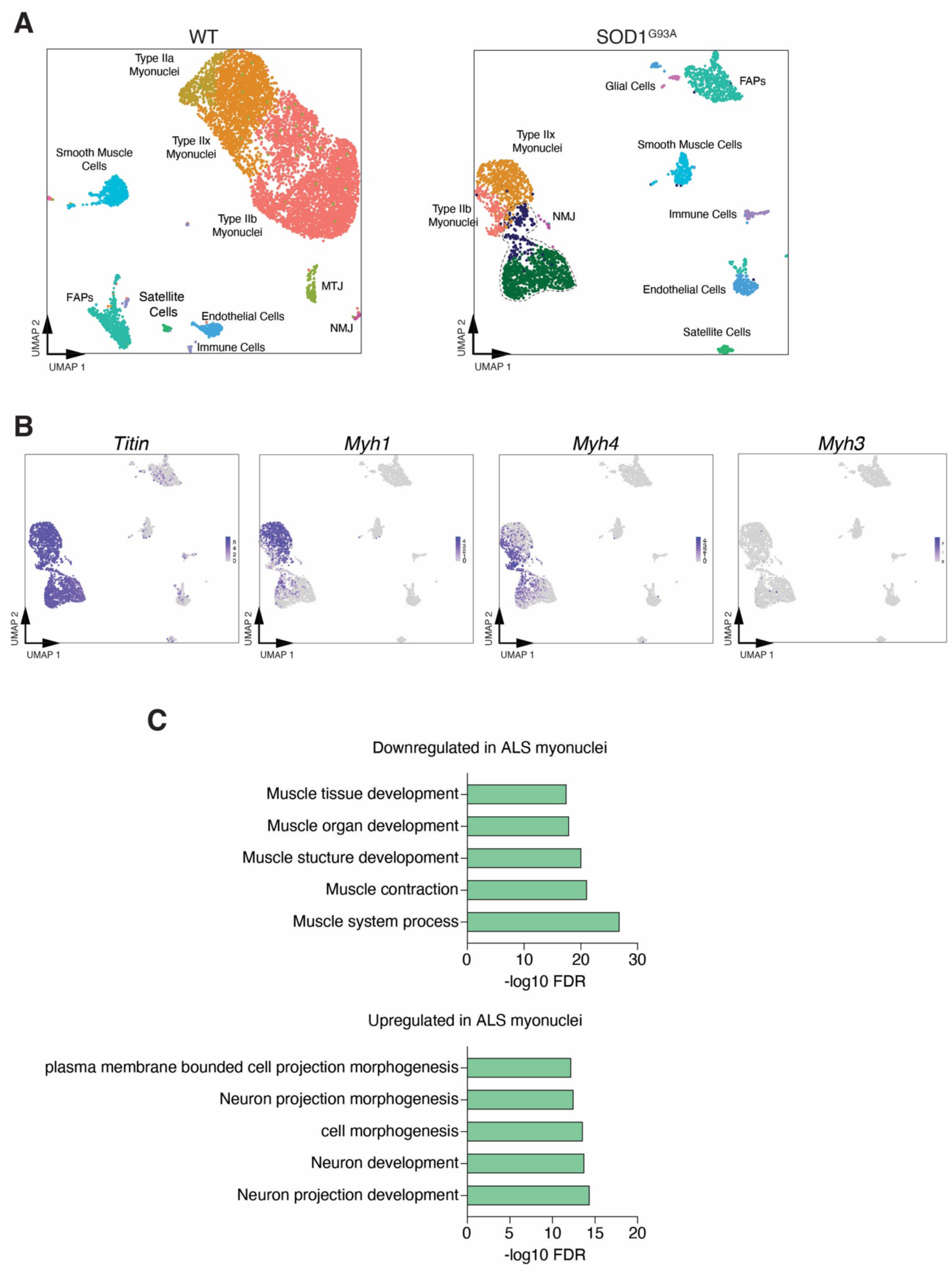
Single nucleus RNA sequencing of the SOD1^693A^ mouse muscle shows a heterogeneous myonuclear transcriptional response. (**A**) UMAP representation shows unbiased clustering of snRNA-seq data from 2 months of age wild-type (WT) or the SOD1^693A^ ALS mice. snRNA-seq was performed on tibialis anterior muscles. (**B**) Feature plots of canonical markers of mature myonuclei in the SOD1^693A^ mouse model. (**C**) Gene Ontology (GO) analysis of the DEGs from ALS myonuclei (Figure 1A), showing significantly changed biological processes with a false discovery rate (FDR) ≤ E-50 criterion. Gene lists used for GO analysis are shown in Supplemental Table 1.

**Supplemental Figure 2.**
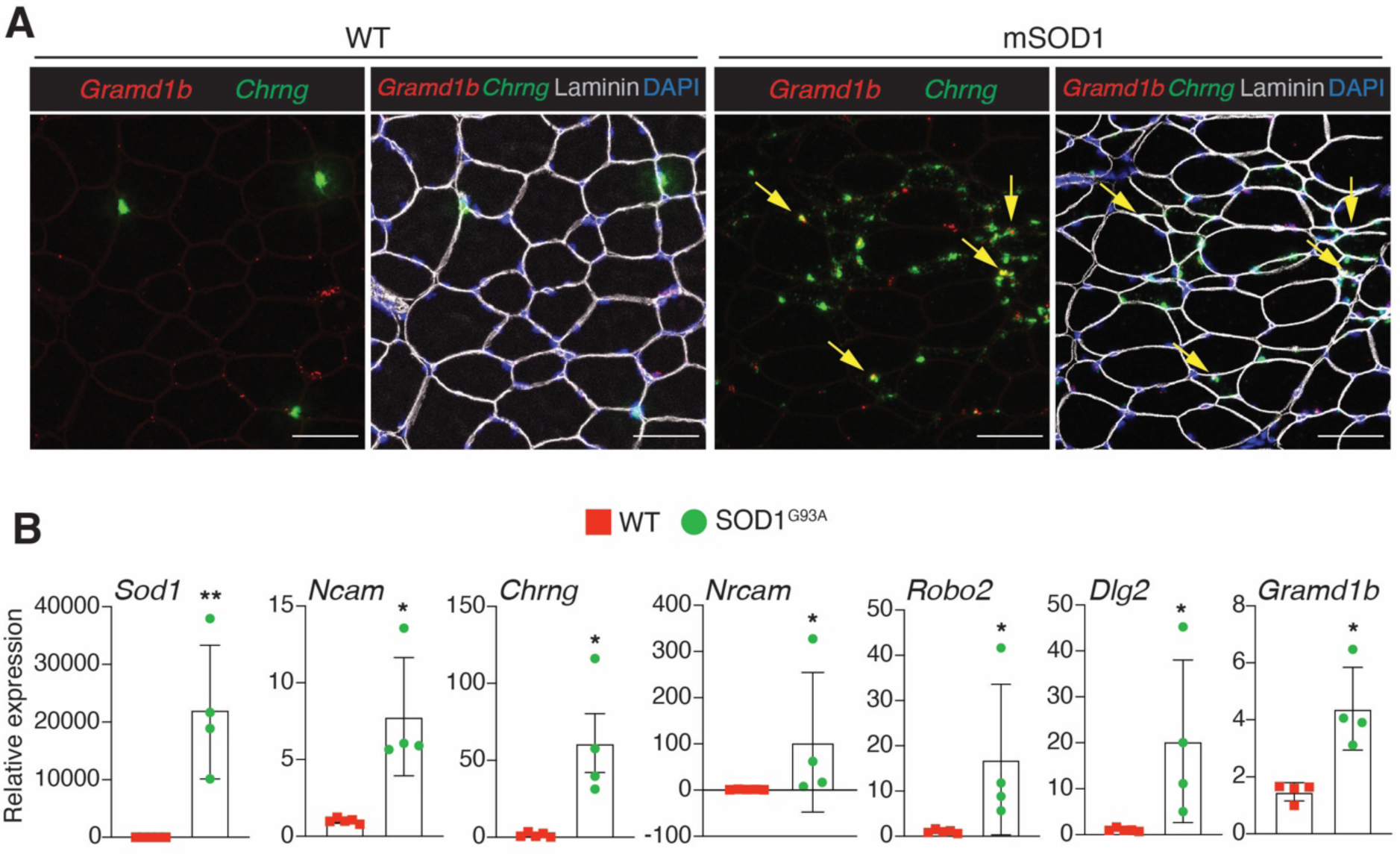
Validation of upregulated genes found in the SOD1^G93A^ mouse snRNA-seq data. (**A**) Representative images of smRNA-FISH for *Chrng* and *Gramd1b* on WT and SOD1^G93A^ TA muscles at 2 months of age. Scale bar, 50μm. (**B**) Reverse transcriptase quantitative PCR (RT-qPCR) analysis for the indicated genes that are upregulated in SOD1^G93A^ TA muscles. Data are presented as mean ± SD. Statistical test used was Mann-Whitney test; *p<0.05, **p<0.01.

**Supplemental Figure 3.**
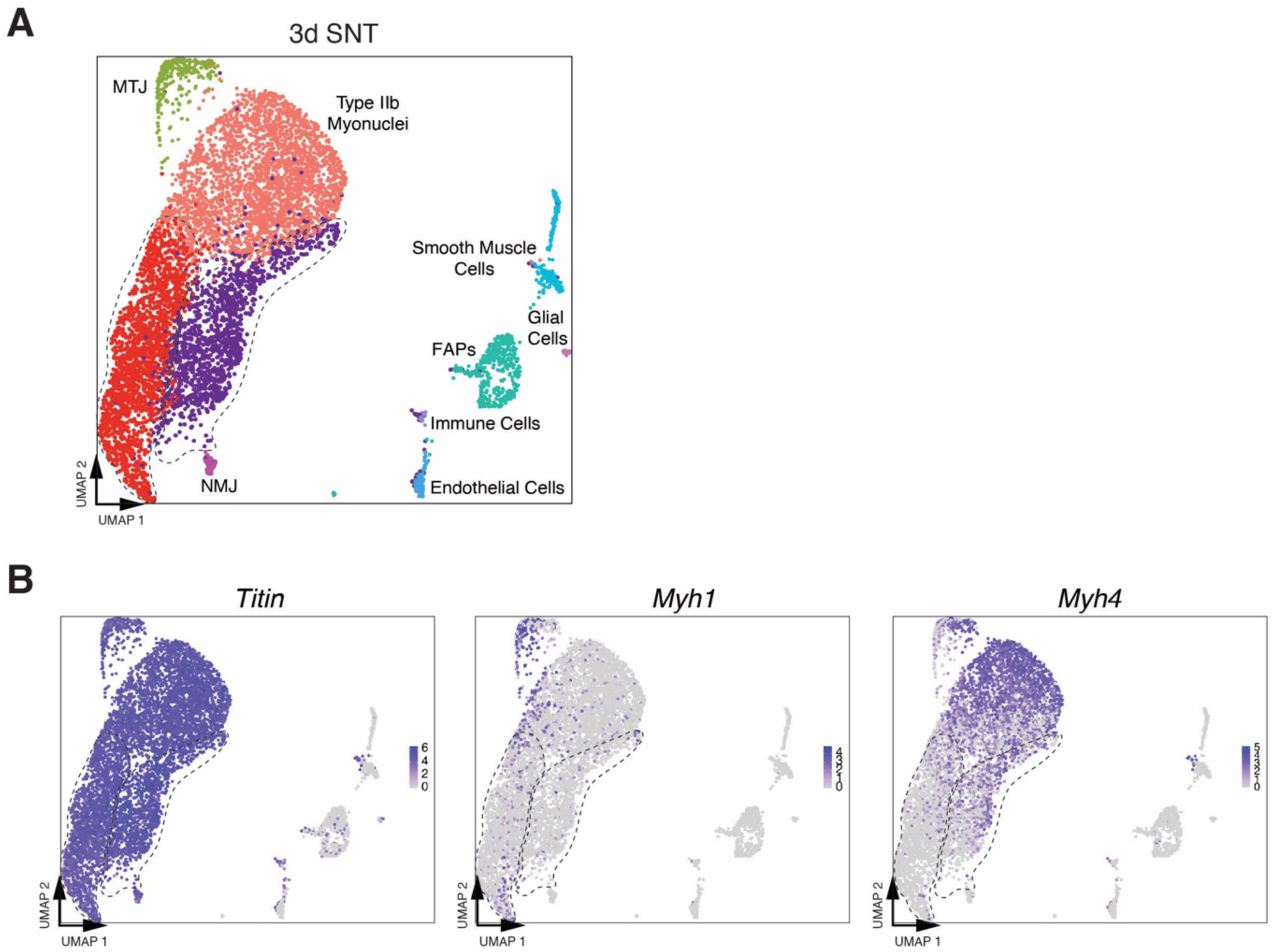
Single nucleus RNA sequencing after 3 days of sciatic nerve transection (3d SNT) shows two denervation-responsive myonuclear populations. (**A**) Unbiased clustering of snRNA-seq data from 3d SNT TA muscle represented on a UMAP. The two denervation-responsive clusters are circled with the dotted lines. (**B**) Feature plots of canonical markers of mature myonuclei in the 3d SNT mouse model.

**Supplemental Figure 4.**
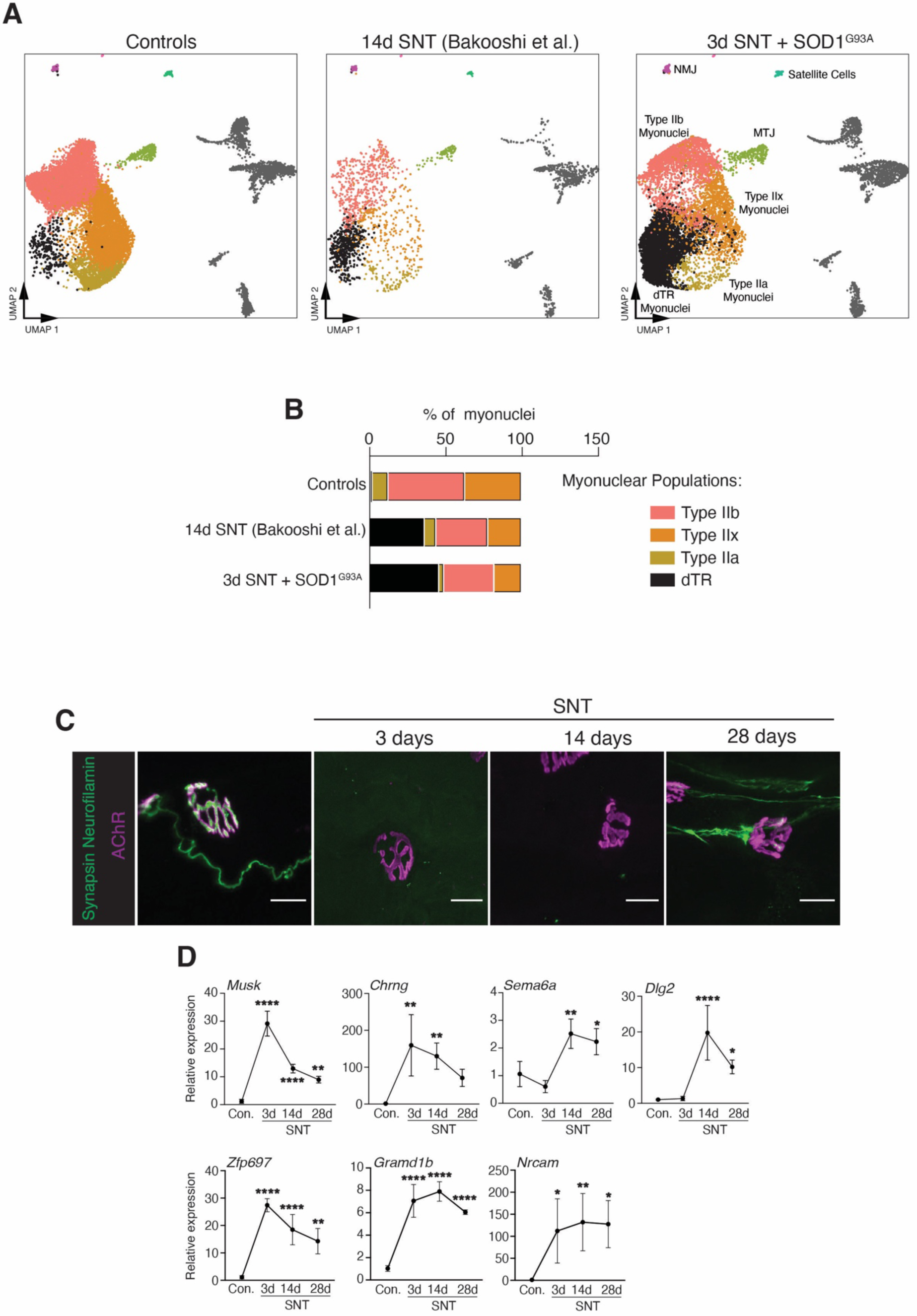
Myonuclear transcriptional response is conserved at later time points following SNT. (**A**) Integrated UMAP split by snRNA-seq data from Controls (Contralateral muscle 14d SNT, 2 months WT and 5 months WT), 14d SNT, and 3d SNT+ SOD1^G93A^ muscles showing the conserved transcriptional signature of the dTR myonuclei cluster. Non-myonuclear populations are colored gray. (**B**) Comparison of the proportions of myonuclear populations excluding NMJ and MTJ between Controls, 14d SNT, and 3d SNT+ SOD1^G93A^. (**C**) Representative staining of AChRs, axons (neurofilament), and nerve terminals (synapsin) from gastrocnemius (GA) muscles from WT adult mice at different time points following SNT. Scale bar, 10μm. (**D**) RT-qPCR analysis showing the dynamic transcriptional response of the indicated genes at different time points following SNT. (control n=5, 3d and 28d SNT n=3; and 17d SNT n=4). Data are presented as mean ± SD. Statistical test used was one-way ANOVA with Tukey’s correction for multiple comparisons; *p<0.05, **p<0.01, ****p<0.0001.

**Supplemental Figure 5.**
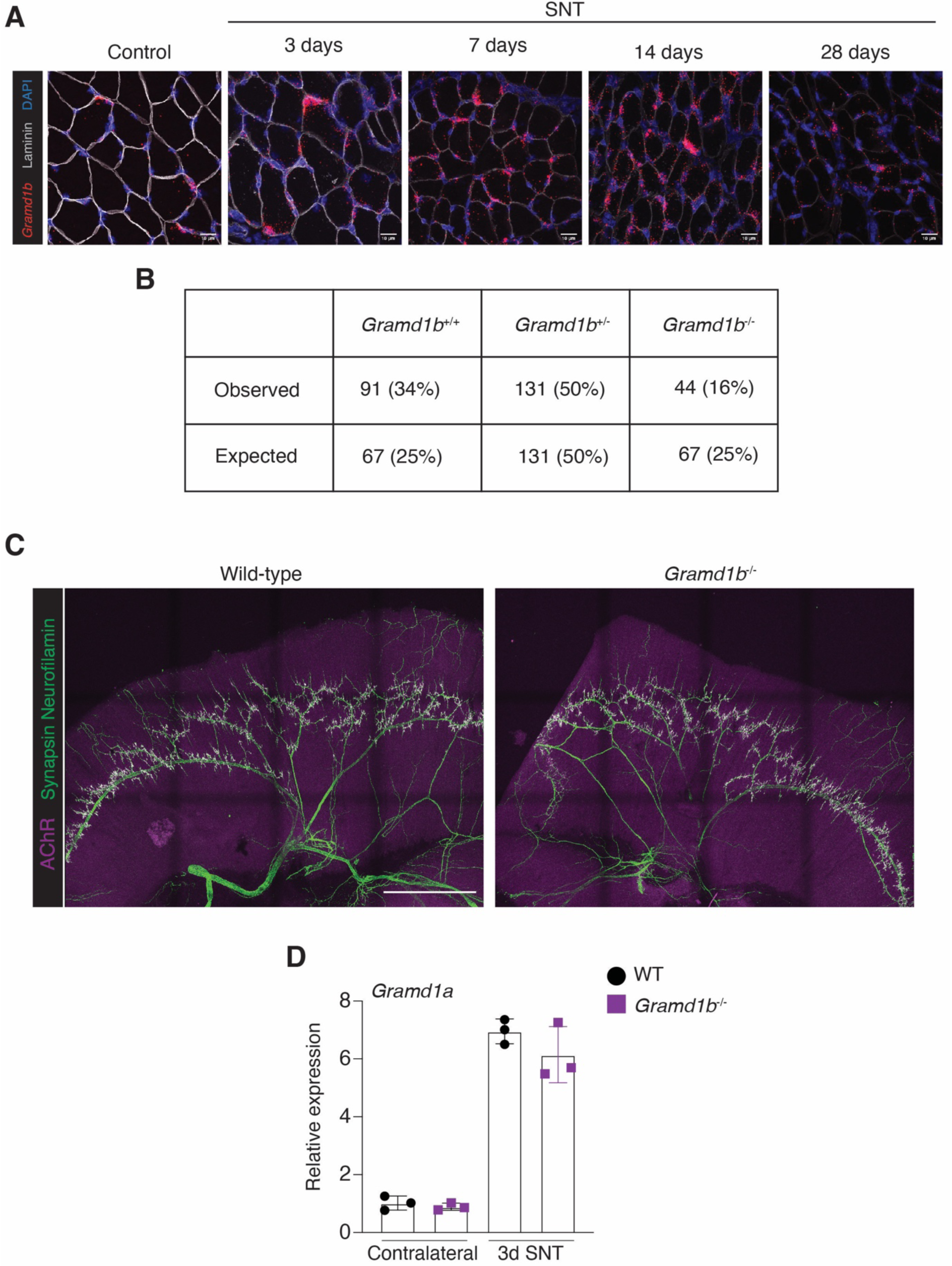
Characterization of *Gramd1b* expression and *Gramd1b*^-/-^ mice. **(A)** Representative images of smRNA-FISH for *Gramd1b* on WT and SNT GA muscles of an adult mouse. Scale bars, 10μm. (**B**) Mendelian ratios of *Gramd1b*^+/+^ (WT), *Gramd1b*^+/-^, and *Gramd1b*^-/-^ mice. Litters were analyzed P7-P10. There is significant reduction of Gramd1b^-/-^ mice (p=0.0149). (**C**) Representative whole-mount diaphragm immunostaining for AChR, axons (neurofilament), and nerve terminals (synapsin) of WT and *Gramd1b*^-/-^ E18 mice. Scale bar, 1 mm. (**D**) RT-qPCR analysis for *Gramd1a* in WT and *Gramd1b*^-/-^ gastrocnemius muscles (contralateral and after 14 days of SNT). Data are presented as mean ± SD. Statistical tests used were (**B**) Chi-square and (**D**) unpaired t-test.

**Supplemental Figure 6.**
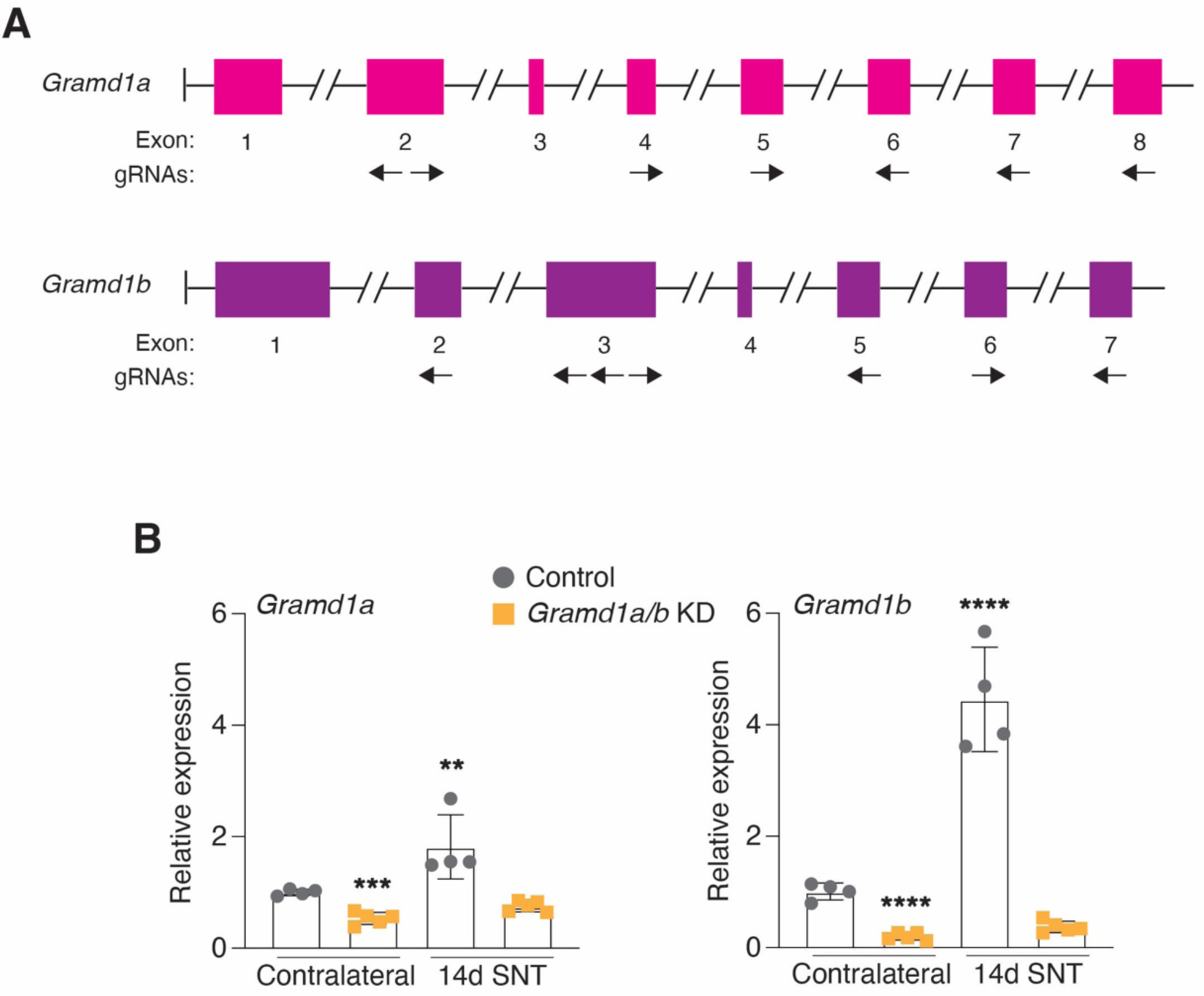
Intramuscular delivery of myoAAV viruses containing gRNAs for *Gramd1a/b* successfully downregulate both genes in contralateral and denervated muscles. (**A**) Schematic representation of sites of gRNAs within *Gramd1a/b* exons. (**B**) *Rosa26*^Cas9^ mice were injected intramuscularly with myoAAVs containing *Gramd1a/b* gRNAs, 14 days after injection, SNT was performed and muscles were collected 14 days after (control n=5, *Gramd1a/b* KD n=4). RT-qPCR analysis of *Gramd1a/b* in EDL muscles. Data are presented as mean ± SD. Statistical test used was unpaired t-test; **p<0.01, ***p<0.001, ****p<0.0001.

## References

1. DeChiara TM, B.D., Valenzuela DM, Simmons MV, Poueymirou WT, Thomas S, Kinetz E, Compton DL, Rojas E, Park JS, Smith C, DiStefano PS, Glass DJ, Burden SJ, Yancopoulos GD, The receptor tyrosine kinase MuSK is required for neuromuscular junction formation in vivo. Cell, 1996. 85(4): p. 501–512.

2. Gautam, M., et al., Defective neuromuscular synaptogenesis in agrin-deficient mutant mice. Cell, 1996. 85(4): p. 525–35.

3. Kim, N., Stiegler, A. L., Cameron, T. O., Hallock, P. T., Gomez, A. M., Huang, J. H., … & Burden, S. J., Lrp4 is a receptor for Agrin and forms a complex with MuSK. Cell, 2008. 135(2): p. 334–342.

4. Okada, K., et al., The muscle protein Dok-7 is essential for neuromuscular synaptogenesis. Science, 2006. 312(5781): p. 1802-5.

5. Weatherbee, S.D., K.V. Anderson, and L.A. Niswander, LDL-receptor-related protein 4 is crucial for formation of the neuromuscular junction. Development, 2006. 133(24): p. 4993–5000.

6. Gautam, M., et al., Failure of postsynaptic specialization to develop at neuromuscular junctions of rapsyn-deficient mice. Nature, 1995. 377(6546): p. 232-6.

7. Tintignac, L.A., H.R. Brenner, and M.A. Ruegg, Mechanisms Regulating Neuromuscular Junction Development and Function and Causes of Muscle Wasting. Physiol Rev, 2015. 95(3): p. 809–52.

8. Clark, J.A., et al., Axonal degeneration, distal collateral branching and neuromuscular junction architecture alterations occur prior to symptom onset in the SOD1(G93A) mouse model of amyotrophic lateral sclerosis. J Chem Neuroanat, 2016. 76(Pt A): p. 35–47.

9. Bermedo-Garcia, F., et al., Functional regeneration of the murine neuromuscular synapse relies on long-lasting morphological adaptations. BMC Biol, 2022. 20(1): p. 158.

10. Jang, Y.C. and H. Van Remmen, Age-associated alterations of the neuromuscular junction. Exp Gerontol, 2011. 46(2-3): p. 193–8.

11. Oda, K., Age changes of motor innervation and acetylcholine receptor distribution on human skeletal muscle fibres. J Neurol Sci, 1984. 66(2-3): p. 327–38.

12. Balice-Gordon, R.J., Age-related changes in neuromuscular innervation. Muscle Nerve Suppl, 1997. 5: p. S83–7.

13. Carlson, B.M., The Biology of Long-Term Denervated Skeletal Muscle. Eur J Transl Myol, 2014. 24(1): p. 3293.

14. Hallock, P.T., et al., Dok-7 regulates neuromuscular synapse formation by recruiting Crk and Crk-L. Genes Dev, 2010. 24(21): p. 2451–61.

15. Bergamin, E., et al., The cytoplasmic adaptor protein Dok7 activates the receptor tyrosine kinase MuSK via dimerization. Mol Cell, 2010. 39(1): p. 100–9.

16. Burden, S.J., R.L. DePalma, and G.S. Gottesman, Crosslinking of proteins in acetylcholine receptor-rich membranes: association between the beta-subunit and the 43 kd subsynaptic protein. Cell, 1983. 35(3 Pt 2): p. 687-92.

17. Zhu, D., W.C. Xiong, and L. Mei, Lipid rafts serve as a signaling platform for nicotinic acetylcholine receptor clustering. J Neurosci, 2006. 26(18): p. 4841–51.

18. Schaeffer, L., A. de Kerchove d’Exaerde, and J.P. Changeux, Targeting transcription to the neuromuscular synapse. Neuron, 2001. 31(1): p. 15–22.

19. Dos Santos, M., et al., Single-nucleus RNA-seq and FISH identify coordinated transcriptional activity in mammalian myofibers. Nat Commun, 2020. 11(1): p. 5102.

20. Petrany, M.J., et al., Single-nucleus RNA-seq identifies transcriptional heterogeneity in multinucleated skeletal myofibers. Nat Commun, 2020. 11(1): p. 6374.

21. Kim, M., et al., Single-nucleus transcriptomics reveals functional compartmentalization in syncytial skeletal muscle cells. Nat Commun, 2020. 11(1): p. 6375.

22. Lai, Y., et al., Multimodal cell atlas of the ageing human skeletal muscle. Nature, 2024. 629(8010): p. 154-164.

23. Bowen, D.C., et al., Localization and regulation of MuSK at the neuromuscular junction. Dev Biol, 1998. 199(2): p. 309–19.

24. Tsay, H.J. and J. Schmidt, Skeletal muscle denervation activates acetylcholine receptor genes. J Cell Biol, 1989. 108(4): p. 1523–6.

25. Kedlian, V.R., et al., Human skeletal muscle aging atlas. Nat Aging, 2024. 4(5): p. 727–744.

26. Witzemann, V., H.R. Brenner, and B. Sakmann, Neural factors regulate AChR subunit mRNAs at rat neuromuscular synapses. J Cell Biol, 1991. 114(1): p. 125–41.

27. Castets, P., et al., mTORC1 and PKB/Akt control the muscle response to denervation by regulating autophagy and HDAC4. Nat Commun, 2019. 10(1): p. 3187.

28. Sandhu, J., et al., Aster Proteins Facilitate Nonvesicular Plasma Membrane to ER Cholesterol Transport in Mammalian Cells. Cell, 2018. 175(2): p. 514–529 e20.

29. Naito, T., et al., Movement of accessible plasma membrane cholesterol by the GRAMD1 lipid transfer protein complex. Elife, 2019. 8.

30. Xiao, X., et al., Hepatic nonvesicular cholesterol transport is critical for systemic lipid homeostasis. Nat Metab, 2023. 5(1): p. 165–181.

31. Xiao, X., et al., Aster-B-dependent estradiol synthesis protects female mice from diet-induced obesity. J Clin Invest, 2024. 134(4).

32. Ferrari, A., et al., Aster Proteins Regulate the Accessible Cholesterol Pool in the Plasma Membrane. Mol Cell Biol, 2020. 40(19).

33. Ferrari, A., et al., Aster-dependent nonvesicular transport facilitates dietary cholesterol uptake. Science, 2023. 382(6671): p. eadf0966.

34. Martineau, E., et al., Dynamic neuromuscular remodeling precedes motor-unit loss in a mouse model of ALS. Elife, 2018. 7.

35. Hegedus, J., et al., Preferential motor unit loss in the SOD1 G93A transgenic mouse model of amyotrophic lateral sclerosis. J Physiol, 2008. 586(14): p. 3337–51.

36. Vinsant, S., et al., Characterization of early pathogenesis in the SOD1(G93A) mouse model of ALS: part II, results and discussion. Brain Behav, 2013. 3(4): p. 431–57.

37. Laraia, L., et al., The cholesterol transfer protein GRAMD1A regulates autophagosome biogenesis. Nat Chem Biol, 2019. 15(7): p. 710–720.

38. Chemello, F., et al., Degenerative and regenerative pathways underlying Duchenne muscular dystrophy revealed by single-nucleus RNA sequencing. Proc Natl Acad Sci U S A, 2020. 117(47): p. 29691–29701.

39. Bakooshli, M.A., et al., Regeneration of neuromuscular synapses after acute and chronic denervation by inhibiting the gerozyme 15-prostaglandin dehydrogenase. Sci Transl Med, 2023. 15(717): p. eadg1485.

40. Bai, L., et al., Motoneurons innervation determines the distinct gene expressions in multinucleated myofibers. Cell Biosci, 2022. 12(1): p. 140.

41. Valenzuela, D.M., et al., Receptor tyrosine kinase specific for the skeletal muscle lineage: expression in embryonic muscle, at the neuromuscular junction, and after injury. Neuron, 1995. 15(3): p. 573–84.

42. Huang, X., J. Jiang, and J. Xu, Denervation-Related Neuromuscular Junction Changes: From Degeneration to Regeneration. Front Mol Neurosci, 2021. 14: p. 810919.

43. Hippenmeyer, S., et al., ETS transcription factor Erm controls subsynaptic gene expression in skeletal muscles. Neuron, 2007. 55(5): p. 726–40.

44. Andersen, J.P., et al., Aster-B coordinates with Arf1 to regulate mitochondrial cholesterol transport. Mol Metab, 2020. 42: p. 101055.

45. Thurkauf, M., et al., Fast, multiplexable and efficient somatic gene deletions in adult mouse skeletal muscle fibers using AAV-CRISPR/Cas9. Nat Commun, 2023. 14(1): p. 6116.

46. Platt, R.J., et al., CRISPR-Cas9 knockin mice for genome editing and cancer modeling. Cell, 2014. 159(2): p. 440–55.

47. Tabebordbar, M., et al., Directed evolution of a family of AAV capsid variants enabling potent muscle-directed gene delivery across species. Cell, 2021. 184(19): p. 4919–4938 e22.

48. Liu, W., et al., Inducible depletion of adult skeletal muscle stem cells impairs the regeneration of neuromuscular junctions. Elife, 2015. 4.

49. Rebolledo, D.L., et al., Denervation-induced skeletal muscle fibrosis is mediated by CTGF/CCN2 independently of TGF-beta. Matrix Biol, 2019. 82: p. 20–37.

50. Lin, H., et al., Decoding the transcriptome of denervated muscle at single-nucleus resolution. J Cachexia Sarcopenia Muscle, 2022. 13(4): p. 2102–2117.

51. Zhang, Y., et al., A molecular pathway for cancer cachexia-induced muscle atrophy revealed at single-nucleus resolution. Cell Rep, 2024. 43(8): p. 114587.

52. Sanes, J.R., L.M. Marshall, and U.J. McMahan, Reinnervation of muscle fiber basal lamina after removal of myofibers. Differentiation of regenerating axons at original synaptic sites. J Cell Biol, 1978. 78(1): p. 176–98.

53. Marshall, L.M., J.R. Sanes, and U.J. McMahan, Reinnervation of original synaptic sites on muscle fiber basement membrane after disruption of the muscle cells. Proc Natl Acad Sci U S A, 1977. 74(7): p. 3073–7.

54. Martyn, J.A., M.J. Fagerlund, and L.I. Eriksson, Basic principles of neuromuscular transmission. Anaesthesia, 2009. 64 **Suppl 1**: p. 1–9.

55. Fröberg, S.O., Determination of muscle lipids. Biochimica et Biophysica Acta (BBA)-Lipids and Lipid Metabolism, 1967. 144(1): p. 14–22.

56. Fischbeck, K.H., E. Bonilla, and D.L. Schotland, Freeze-fracture analysis of plasma membrane cholesterol in fast- and slow-twitch muscles. J Ultrastruct Res, 1982. 81(1): p. 117–23.

57. Williams, K.D. and D.O. Smith, Cholesterol conservation in skeletal muscle associated with age- and denervation-related atrophy. Brain Res, 1989. 493(1): p. 14–22.

58. Ikonen, E., Cellular cholesterol trafficking and compartmentalization. Nat Rev Mol Cell Biol, 2008. 9(2): p. 125–38.

59. Ikonen, E. and X. Zhou, Cholesterol transport between cellular membranes: A balancing act between interconnected lipid fluxes. Dev Cell, 2021. 56(10): p. 1430–1436.

60. Luo, J., H. Yang, and B.L. Song, Mechanisms and regulation of cholesterol homeostasis. Nat Rev Mol Cell Biol, 2020. 21(4): p. 225–245.

61. Gurney, M.E., et al., Motor neuron degeneration in mice that express a human Cu,Zn superoxide dismutase mutation. Science, 1994. 264(5166): p. 1772-5.

62. Concordet, J.P. and M. Haeussler, CRISPOR: intuitive guide selection for CRISPR/Cas9 genome editing experiments and screens. Nucleic Acids Res, 2018. 46(W1): p. W242–W245.

63. Sakuma, T., et al., Multiplex genome engineering in human cells using all-in-one CRISPR/Cas9 vector system. Sci Rep, 2014. 4: p. 5400.

64. Challis, R.C., et al., Systemic AAV vectors for widespread and targeted gene delivery in rodents. Nat Protoc, 2019. 14(2): p. 379–414.

65. Kann, A.P. and R.S. Krauss, Multiplexed RNAscope and immunofluorescence on whole-mount skeletal myofibers and their associated stem cells. Development, 2019. 146(20).

66. Fleming, S.J., et al., Unsupervised removal of systematic background noise from droplet-based single-cell experiments using CellBender. Nat Methods, 2023. 20(9): p. 1323–1335.

67. Hao, Y., et al., Dictionary learning for integrative, multimodal and scalable single-cell analysis. Nat Biotechnol, 2024. 42(2): p. 293–304.

68. Bernstein, N.J., et al., Solo: Doublet Identification in Single-Cell RNA-Seq via Semi-Supervised Deep Learning. Cell Syst, 2020. 11(1): p. 95–101 e5.

69. Vandesompele, J., et al., Accurate normalization of real-time quantitative RT-PCR data by geometric averaging of multiple internal control genes. Genome Biol, 2002. 3(7).

70. Chen, J., et al., ToppGene Suite for gene list enrichment analysis and candidate gene prioritization. Nucleic acids research, 2009. 37(suppl_2): p. W305-W311.

71. Kemp, T.J., et al., Identification of Ankrd2, a novel skeletal muscle gene coding for a stretch-responsive ankyrin-repeat protein. Genomics, 2000. 66(3): p. 229–41.

72. Ravindra Kumar, S., et al., Multiplexed Cre-dependent selection yields systemic AAVs for targeting distinct brain cell types. Nat Methods, 2020. 17(5): p. 541–550.

